# A Proteome-scale Analysis of Vertebrate Protein Amino Acid Occurrence: Thermoadaptation shows a Correlation with Protein Solvation but less so with Dynamics

**DOI:** 10.1101/2020.08.10.244558

**Authors:** Z.L. Li, M Buck

## Abstract

Despite differences in behaviors and living conditions, vertebrate organisms share the great majority of proteins, often with subtle differences in amino acid sequence. Here, we present a simple way to analyze the difference in amino acid occurrence by comparing highly homologous proteins on a sub-proteome level between several vertebrate model organisms. Specifically, we use this method to identify a pattern of amino acid conservation as well as a shift in amino acid occurrence between homeotherms (warm-blooded species) and poikilotherms (cold-blooded species). Importantly, this general analysis and a specific example further establish a correlation, if not likely connection between the thermoadaptation of protein sequences and two of their physical features: a possible change in their protein dynamics and, even more strongly, in their solvation. For poikilotherms, such as frog and fish, the lower body temperature is expected to increase the association of proteins due to a decrease in protein internal dynamics. In order to prevent overly-sticky protein association at low temperatures, the use of amino acids suggests that poikilotherms enhance the solvation of their proteins by favoring polar groups on their protein’s surface. This feature appears to dominate over possible changes in dynamics. The results suggest that a general trend for amino acid choice is part of the mechanism for thermoadaptation of vertebrate organisms at the molecular level.

## Introduction

The diversity of living organisms is a result of the evolution of genes with respect to their encoded biological macromolecules. In general, todays organisms share a large number of homologous proteins of similar sequences and function. Amino acid sequences evolved in a way to optimize their protein’s function, allowing the organisms to adapt to their different environments. The role of site specific mutations has been demonstrated in multiple proteins, as an organism adapts to extreme cold (e.g. L-Lactate dehydrogenase A chain, LDHA),^1^ oxygen availability (Hemoglobin),^2^ light (Rhodopsin),^3,4^ high altitude (p53),^5^ or the scarcity of food (e.g. Melanocortin 4 receptor).^6^ Comparing the proteome of two organisms at the level of amino acid occurrence in highly homologous proteins may provide a clue as to how they may have adapted to different living conditions. Yet to our knowledge, such analyses have not been carried out for higher level organisms, specifically for vertebrate model systems. Here we present a simple analysis, which reveals changes in amino acid utilization and attempt to rationalize the results in the context of cold adaptation in vertebrates.

In spite of sharing a large number of near identical proteins between the different organisms, vertebrates have an obvious and wide ranging difference in body temperature, most noticeably between warm-blooded and cold-blooded animals (homeotherms and poikilotherms, respectively). The activity of proteins is typically highly sensitive to temperatures. In a human being, a small increase of body temperature by 3-5 °C can seriously damage cells, tissues and organs. Thus, it is remarkable that similar proteins could adapt and function at a broad range of temperatures. A cold-blooded organism, for example the African clawed frog, has a wide tolerance for changes in body temperature ranging from 10 to 30°C; for zebrafish, a range of body temperature between 16°C and 30°C allows it to swim.^7^ Both are lower than the temperature of humans (median 37°C), other mammals (median 38°C) and birds (median 40°C). The apparent drop in body temperature for cold-blooded organisms would generally attenuate, or “quench” many protein fluctuations, which – based on entropy considerations – should lead to enhanced protein association. This is usually detrimental to the efficiency of enzyme catalysis or protein signaling processes.

In order to adjust to the environmental temperature that alters protein dynamics and function, an organism would have accumulated certain types of amino acid changes (mutations), possibly at preferred sites, as part of a process of adaptation to a changed body temperature. For cold-blooded organisms, those changes may alter the protein internal dynamics and more likely the affinity of the protein for water. The adaptation of a protein surface, but also in some cases of interior positions, to water represents an important factor influencing protein recognition and activity.^8–10^ The “life on the edge of solubility” hypothesis posits that the expression of a protein in the cellular environment occurs very close to its solubility limit under most physiological solution.^11^ Therefore, even few mutations at certain sites which alter protein solvation may, in some cases dramatically influence the total protein solubility (e.g. sickle cell Hemoglobin). Overall, the dynamics and solubility of a protein are both crucial for its activity and for the well-being of the organism.^11–22^ Mechanisms of protein thermoadaptation related to protein stability and solvation at high temperatures have been recognized in several proteins studied, e.g. by comparing a protein from a normal and a thermophilic setting. ^1,23,24^. However, while investigations on individual, usually prokaryotic organisms have led to pattern of change in amino acid occurrence, it is unknown whether a similar general principle for thermoadaptation may exist at a proteome-wide level of scale for higher level organisms. This prompted us to investigate a change in amino acid occurrence with respect to the role of key amino acids in protein dynamics and solvation in the thermoadaptation of proteins by analyzing sub-proteomes of multiple vertebrate organisms.

## Results

### Conservation and Shift of Occurrence of 20 Amino Acids Over Sub-proteomes of Model Vertebrate Organisms

We compared the sequence of proteins with the same protein name in Uniprot and <5% difference in amino acid sequence length between human (Homo sapiens) and several organisms, typically chosen as representative models or as a reference in proteome studies of vertebrates (Fig. 1a): Compared with human proteins are those of chimpanzee (Pan troglodytes), mouse (Mus musculus), chicken (Gallus gallus), African clawed frog (Xenopus laevis), and zebrafish (Danio rerio). This involves 3607, 14046, 1921, 1565, and 2353 protein pairs for the respective human-other organism comparison, fulfilling the simple two criteria of name identity and near identical sequence length (see also Table S1). The sub-proteome comparisons, which result with these two limiting parameters, have more than 98% of the proteins paired with greater 30% sequence identity and can be considered homologous. For each protein -a homolog pair between human and the chosen organism-we compared the difference in the content for each one of the 20 amino acids (scaling by protein lengths) to give the value for a shift in occurrence, if any, for that amino acid ΔC_i_(AA) (see Methods). For example, a 143 amino acid residue sequence constitutes the Hemoglobin subunit alpha (HSA) in zebrafish, of which 18 positions are alanine. The corresponding human protein has 21 alanines in 142 amino acids. Thus, the difference in the content of alanine in Hemoglobin subunit alpha between zebrafish (12.59%) and human (14.79%) is −2.20 %. In the same way, the differences were calculated for the 2353 paired proteins between zebrafish and human. The % difference for Ala is collected into 9 bins to give an overview of the shifts in occurrence for all protein pairs and then plotted as a histogram (Fig. 1b). Rather than using the number of protein pairs in each bin as the y-axis we can divide by their number to give the % of proteins in that bin. Statistically, of the 2353 proteins, 220 proteins (9%) have only subtle alternations in the sequence content of alanine within ±0.125%. For 823 proteins (35%), zebrafish has less alanine than the human protein with a difference in the absolute content of ≤ −1%. Similarly, there are 444 proteins (19%) in zebrafish which have more alanine ≥ 1% compared to the human protein homolog. Therefore, there is an obvious asymmetry or shift in amino acid use of alanine over a considerable proportion of the proteome between zebrafish and human proteins. In the same way, the results for all the other amino acids between the above-mentioned model organisms are analyzed and summarized in Fig. 1c (also Fig. S1a).

**Figure 1:**
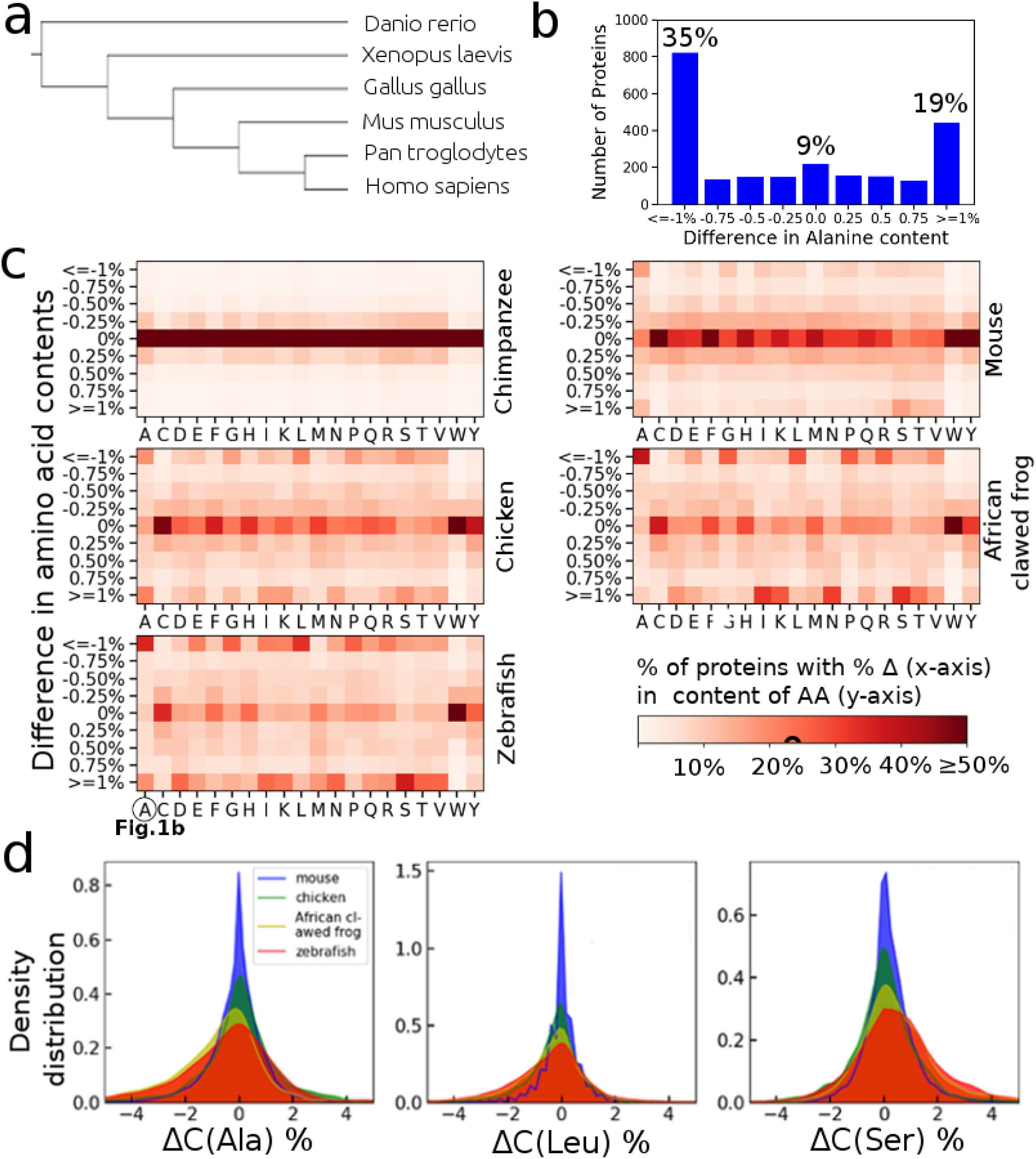
Conservation and change of the occurrence of the 20 amino acids, comparing paired sub-proteomes of model vertebrate organisms. (a) Phylogenetic tree of representative organisms considered in this study: Human (Homo sapiens, chimpanzee (Pan troglodytes), mouse (Mus musculus), chicken (Gallus gallus), African clawed frog (Xenopus laevis), and zebrafish (Danio rerio). (b) The distribution of C(Ala) for 2353 pair-wise proteins between zebrafish and human. The difference in the occurrence of alanine of an individual protein pair, Ci(Ala) is put into one of 9 bins from −1% to 1% with an increment of 0.25%. Differences < −1% or >1% are added to the lowest and highest bin, respectively. This is done for all protein pairs, leading to the histogram plotted for Ala, as shown. The distribution of C for the other amino acids and other model organisms are given over the protein pairs in (c). The y-axis value is the extent of the difference in absolute amino acid content (bin-ranges, same as x-axis in b). The x-axis indicates the 20 amino acids. [A(la) is circled for zebrafish as this column corresponds to Fig. 1b]. The color bar indicates the fraction (%) of proteins in the comparison that have a content difference in the bins (see the unscaled y-axis in b and fractions given in figure). (d) Density distribution of C for alanine, leucine and serine between a model organism and human (data for chimpanzee proteins are nearly identical to human and data isnot shown for better visualization). Note that except for the extremes, the binned range is −1 to +1%, showing that a significant number of human/zebrafish compared proteins are in the extreme bins in the figure panels b-d). The statistical significance (p-value) is given in Table S2.

A statistical analysis of the average content of an amino acid in the proteins of a model organism, as well as the difference comparing to the human homolog is given in Table S2 (in terms of p-values). Based on Fig. 1c and Table S2, in general, the extent of variation in amino acid utilization reflect the closeness of a model organism to the human sequences (see Discussion). For example, amino acid sequences between chimpanzee and human proteins are highly identical with only very few changes. Mouse and chicken proteins have moderate changes compared to their human homologs. Remarkably, we noticed a significant directional difference between cold-blooded animals (African clawed frog and zebrafish) and warm-blooded organisms (human, chimpanzee, mouse and chicken), with respect to the extent of variation in several polar/non-polar amino acids.

In the cold-warm blooded organism comparison, the paired protein sequence analysis yields four major observations: i) <u>Phenylalanine (F)</u>, <u>histidine (H)</u>, <u>glutamine (Q)</u>, <u>cysteine</u> (C) and <u>valine (V)</u> experience a relatively small change in their occurrence in the proteins. The average content of <u>tyrosine</u> (Y) and <u>tryptophan</u> (W) in proteins is low, but there is also little change in the occurrence of these two amino acids. ii) Both African clawed frog and zebrafish have a lower negative charge per protein (−0.57 and −0.94 *e* on average) than the corresponding human proteins. This is mainly achieved by a higher utilization of aspartic acid (D) (see also Fig. S2). The occurrence of <u>glutamic acid</u> (E), <u>lysine</u> (K) and <u>arginine</u> (R) also changes considerably, but mainly in a way of mutual exchange of D with E, or of K with R. iii) A great number of human proteins have a higher content of <u>alanine (A)</u>, <u>glycine (G)</u>, <u>leucine (L)</u> and <u>proline (P)</u>, but a lower content of <u>isoleucine (I)</u> (Fig. 1c, d, and Table S2). It is generally accepted that such changes may influence the dynamics of the polypeptide chain, with glycine, alanine, providing greater flexibility whereas isoleucine reduce flexibility due to β-branched sidechains (a prediction we test for two examples below). iv) By contrast, there is an increase in the relative proportion (again in terms of occurrence in the population of paired proteins) of polar amino acids <u>serine (S)</u>, <u>threonine (T)</u>, <u>asparagine (N)</u> as well as <u>methionine (M)</u> in African clawed frog and zebrafish (Fig. 1c, d and Table S2) compared to the warm-blooded organisms. Again, it is thought that this group of amino acids relates to protein solvation (solubility).

For simplicity, we further classified (G), (L, A, I), (K, R), (D, E), (N, S, T, M) into five groups separately and perform the statistical analysis over the combined contents of amino acid in each group. Table 1 shows a directional and sizeable difference between human and African clawed frog/zebrafish in the utilization of glycine, polar versus non-polar amino acids. On average, the absolute content of Glycine (G) and hydrophobic amino acids (L, A, I) are reduced by 0.4% and 0.7% respectively for African clawed frog, and by 0.3% and 0.9% respectively for zebrafish. By contrast, the content of polar amino acids (N, S, T) and methionine (M, actually having polar side chain) are increased, on average in the proteins considered by 1.3% for African clawed frog and by 1.2% for zebrafish. Similar patterns are observed when comparing zebrafish and frog to chicken instead of human proteins, but the extent is reduced (Fig. S1b, Table S1b and Table S3). The utilization of Gly (in chicken) is reduced by 0.3% and 0.2% in going to African clawed frog and zebrafish respectively, while the content of grouped (N, S, T, M) is increased by 1.0% and 1.11%. The content of (L, A, I) are reduced by 1.65% and 0.81% respectively. Overall, these changes suggest a significant difference of utilization of polar and non-polar amino acids between cold-blooded organisms and warm-blooded organisms.

**Table 1:** Statistical analysis of contents of amino acids grouped into 5 types between human and a model organism of (a) mouse, (b) chicken, (c) African clawed frog and (d) zebrafish. 14046, 1921, 1565, and 2353 paired proteins are compared for the analysis, same as Fig.1. The difference in absolute content and the significance (p-value) are added under each column.

|  | Organism | G | A/I/L | K/R | D/E | N/S/T/M |
| --- | --- | --- | --- | --- | --- | --- |
| N=14046 | mouse | 6.6% | 21.7% | 11.49% | 11.6% | 18.9% |
|  | Human | 6.7% | 21.9% | 11.51% | 11.6% | 18.7% |
| | Difference $\Delta$ | **-0.1%** | **-0.2%** | **-0.02%** | **0%** | **0.2%** |
|  | p-value | <0.0001 | <0.0001 | <0.0001 | 0.7383 | <0.0001 |
| N=1921 | chicken | 6.3% | 21.75% | 11.9% | 12.14% | 18.9% |
|  | Human | 6.4% | 21.81% | 11.7% | 12.07% | 18.7% |
| | Difference $\Delta$ | **-0.1%** | **-0.06%** | **0.2%** | **0.07%** | **0.1%** |
|  | p-value | 0.0002 | 0.1552 | <0.0001 | 0.0024 | 0.0102 |
| N=1565 | African Clawed Frog | 5.9% | 21.3% | 11.77% | 12.2% | 19.8% |
|  | Human | 6.4% | 22.0% | 11.76% | 12.1% | 18.5% |
| | Difference $\Delta$ | **-0.4%** | **-0.7%** | **0.01%** | **0.2%** | **1.3%** |
|  | p-value | <0.0001 | <0.0001 | 0.7357 | <0.0001 | <0.0001 |
| N=2353 | zebrafish | 6.2% | 21.5% | 11.61% | 12.0% | 19.5% |
|  | Human | 6.5% | 22.4% | 11.65% | 11.8% | 18.3% |
| | Difference $\Delta$ | **-0.3%** | **-0.9%** | **-0.04%** | **0.2%** | **1.2%** |
|  | p-value | <0.0001 | <0.0001 | 0.1065 | <0.0001 | <0.0001 |

In absence of sequence adaptation, the apparent drop in body temperature for cold-blooded organisms would generally lead to decreased protein internal as well as global fluctuations (i.e. lower the entropy of a proteins), typically this is expected to result in an enhancement in protein association and usually a reduced protein activity. By making the protein even less dynamic (a change of Gly and Ala to other amino acids in going from hot to cold-blooded organisms) the proteins should become even more sticky, thus presenting us with a conundrum to rationalize such changes. Thus, next, we examined this issue specifically with respect to estimates for these two protein features, dynamics and solvation for the cold and warm-blooded model organisms.

### Entropy and Solvation Energy Over Sub-proteomes of Human and Model Organisms

We performed a sequence-based prediction of the polypeptide chain entropy and solvation free energy for the sets of protein homologous pairs examined in Fig. 1. These calculations are based on available experimental values of amino acid backbone entropy and solvation free energy for model peptides, chemical groups, respectively (Fig. S3).^25,26^ We found that African clawed frog and zebrafish proteins have an overall lower backbone entropy in comparison to warm-blooded species, chicken, mouse, chimpanzee and human (Fig. 2a). These results reflect our observation above that frog and fish proteins did not accumulate residues that are expected to increase protein dynamics. Specifically, the proteins of the cold-blooded organisms have a relatively higher abundance of Ile and Val (lowest backbone entropy) and lower content of Ala and Gly (highest backbone entropy). Furthermore, based on the sequence analysis the fish and frog proteins are predicted to have an increased affinity toward water (−100.9 cal mol^−1^ res^−1^ and −144.5 cal mol^−1^ res^−1^ respectively) (Fig. 2a). This is in accord with the increased occurrence of polar amino acids (Ser, Thr, Asn) in cold-blooded organisms. However, even when we consider an estimation of the proteins entropy only accounting for the mainchain an underestimate, importantly, the predicted unfavorable entropy change is much smaller than the increase in solvation. A table showing which amino acids are changed to which on going from human to zebrafish proteins, Fig. 2b shows that Ala and Gly are frequently changed to polar amino acids (Ser, Thr) in cold-blooded organisms. Also, Leucines in human proteins are frequently replaced by Methionine, which is polar near its sidechain terminal.

**Figure 2:**
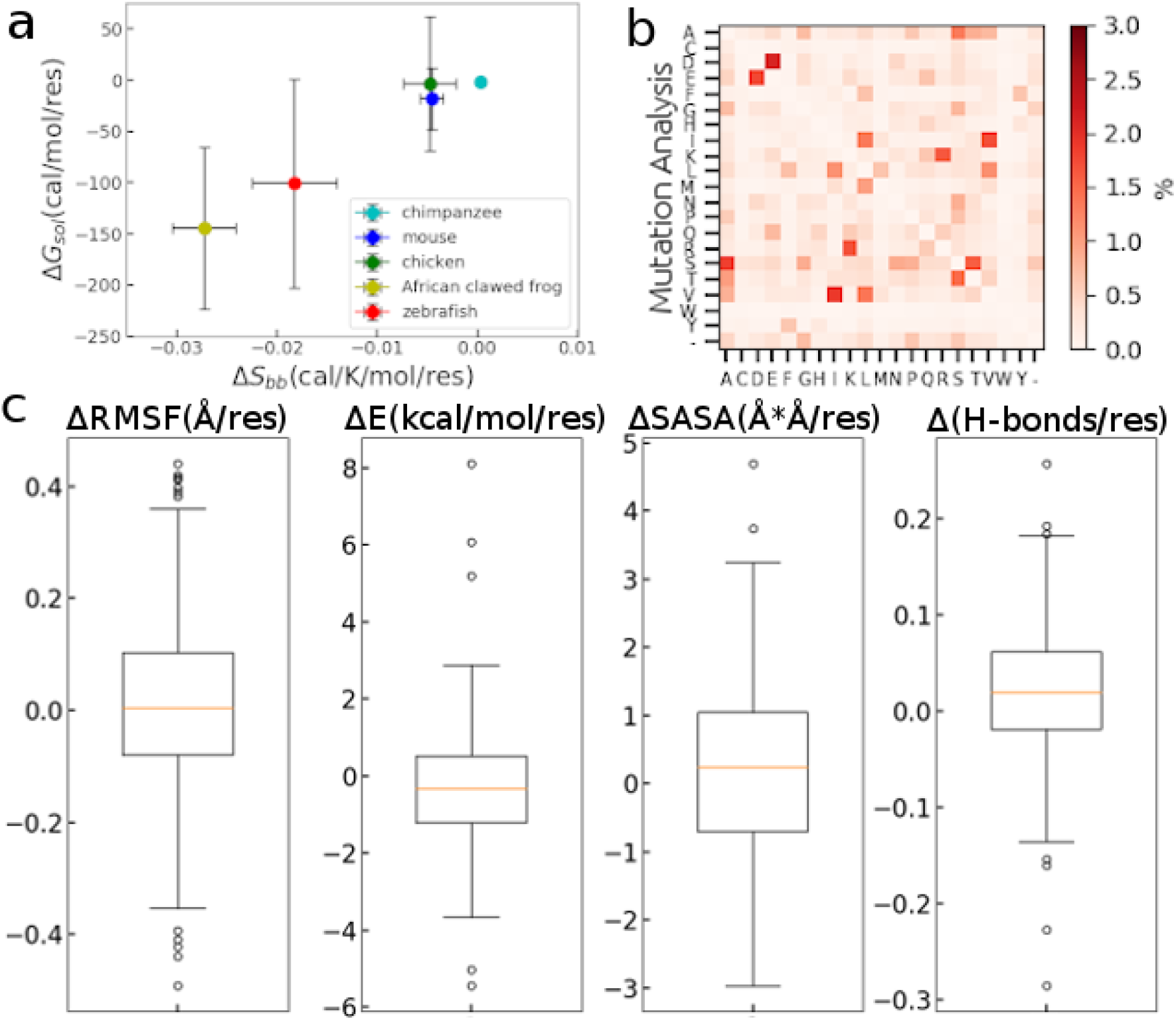
Energetics and entropy analysis over a sub-proteome between human and model organisms. (a) Estimation of difference in entropy and solvation free energy between paired proteins from a model organism and humans. The entropy and solvation energy is averaged over the sequence length of a protein (per amino acid). The difference in average values and the significance (p-value) are as below (chimpanzee, mouse, chicken, African clawed frog, zebrafish versus human): Entropy [cal/K/mol], (0.0003, p=0.128), (−0.0046, p<0.0001), (−0.0048, p<0.0001), (−0.0273, p<0.0001), (−0.0183, p<0.0001); Solvation Energy [cal/mol], (−1.7, p=0.0596), (−18.2, p<0.0001), (−3.5, p=0.547), (−144.5, p<0.0001) (−100.9, p<0.0001). (b) Amino acid mutation analysis for 2353 protein pairs between zebrafish and human. The 21th. row/column indicates amino acid deletion/addition. (c) Energetic analysis for the simulations of 166 paired proteins between human and zebrafish. Boxplots of RMSF (Root mean squared fluctuation), E (Poisson Boltzmann electrostatic interaction), SASA (Solvent available surface area), and H-bonds (number of hydrogen bonds between solvent and protein). All values in e) are scaled by the sequence length, i.e. are the average change per amino acid.

In order to link the amino acid changes to structural and dynamics changes in the proteins with known structures, we carried out a molecular dynamics study on 166 human proteins and their zebrafish homology models.

We noted that 96.4% of the mutations are found at the water-interacting protein surface (0-4 to water); 3.6% are essentially in the interior of the protein (>4). In the 100 ns simulations for the 166 proteins pairs we observed little difference in backbone fluctuations (RMSF of Cα atoms, ΔRMSF= −0.01 Å on average, p=0.9043) between human and zebrafish proteins (Fig. 2c). However, compared to their human homologs, the zebrafish proteins have enhanced Poisson Boltzmann electrostatic interactions with the solvent (ΔE= − 0.32 kcal mol^−1^ res^−1^ on average, p=0.0149) as well as a larger solvent accessible surface area (ΔSASA=0.22 Å^2^ res^−1^ on average, p=0.0333) (Fig. 2c). The protein’s enhanced affinity for water is partially caused by the enhanced enthalpy for hydrogen bond formation at a lower temperature, but since the simulations are carried out at the same temperature, only the difference in sequence, i.e. surface structure contributes. Thus number of protein polar group-water hydrogen bonds is increased ( H-bonds= 0.023 res^−1^ on average, p=0.0003) on going from human to zebrafish proteins (Fig. 2c). These numbers look very small but when multiplied by the average length of a protein, say at 200 residues, 4.6 new hydrogen bonds between the protein and water were formed.

Together, both our sequence-based and structure-based energy analyses support the finding that the zebrafish proteins have an overall enhanced degree of solvation. The solvation increase would lead to a bigger loss of protein solvation energy in protein association.^27^ As discussed below, this may help to prevent an over-sticky association caused by low temperature during enzyme catalysis or cellular signaling processes.

### Dynamics and Solvation in L-Lactate dehydrogenase A chain (LDHA) for Antarctic Fish in Extreme Cold

Lastly, we present a case study of one protein L-Lactate dehydrogenase A chain (LDHA) of Antarctic fishes (scientific name, notothenioid). In this situation, the protein needs to adapt to an extreme cold of −2 to 0 °C. The protein was found to have a decreased substrate binding affinity but an increased rate of catalysis at nearzero temperature.^1^ Looking at the sequence, the LDHA of Antarctic fishes (nothothenioidei group) have an occurrence of amino acids which is consistent with the he sub-proteome level of zebrafish and frog (poikilotherm, or commonly cold-blooded, but living at moderate temperatures),, i.e., the higher abundance of Ser, Thr, Met, Val, and lower content of Ala and Leu (Fig. 3a). Intriguingly, the Antarctic fishes have more Gly residues, opposite from the general direction of the shift of Gly occurrence noted in the two other model organisms, zebrafish and frog, above (Fig. 1 and Fig. 2).

**Figure 3:**
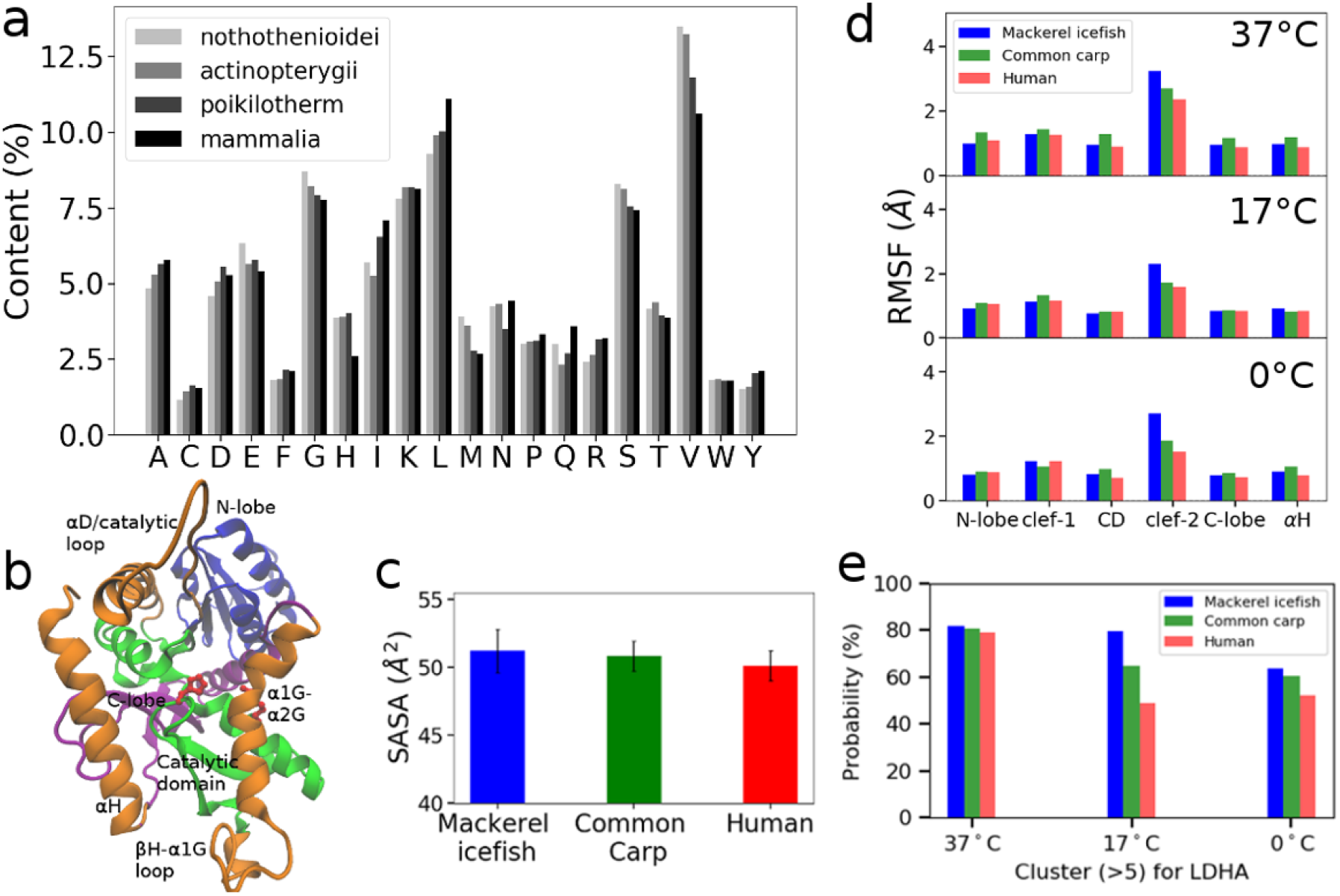
Cold adaptation of L-Lactate dehydrogenase A chain (LDHA) of Antarctic fishes in extreme cold. (a) Comparing the occurrence of different amino acids for Antarctic fishes (nothothenioidei), other moderate temperature environment fishes (actinopterygii), cold-blooded organisms excluding fishes (poikilotherm) and mammalia. (b) LDHA (Mackerel icefish) structures and domain partitioning. N-lobe: 1-94 (blue); Catalytic subdomain: 129-210 (green); C-lobe: 245 to 307 (purple). Three secondary structural elements encompassing the active site are marked in green: D/catalytic loop (95-128); H – 1G loop and 1G-2G (211-244); and H (308 to 331). His 192 and Arg167 are space-filled to indicate the location of the active site. The starting residues of 1-19 are excluded for the analysis due to high flexibility. (c) Surface accessible surface area of LDHA per residue of three different organisms. The difference per residue and significance (p-value) are (1.08 Å^2^, p<0.0001), (0.70 Å^2^, p<0.0001) respectively comparing Mackerel icefish and common carp with human. (d) RMSF (average over amino acid residues in 6 select regions/subdomains) of LDHA of three different organisms at three different temperatures (37, 17 and 0). (e) Cluster analysis of LDHA from three different organisms at different simulation temperatures.

Next, we compared the dynamics of a monomeric LDHA of three different species with available protein structures: Mackerel icefish (a notothenioid fish), living in waters of −2 to 0 °C, common carp living in normal water (ranging from 3 to 35 °C) and lastly Human (37 °C) (Fig. S4a). Molecular dynamics simulations for LDHA proteins of human, common carp and Mackerel icefish were performed in triplicate for 500 ns at three different temperatures: 310K (37), 290K (17) and 273K (0). The LDHA protein is composed of several different sub-domains or lobes: briefly, the N-terminal-, catalytic and the C-terminal lobes (Fig. 3b) and three regions encompassing the active site as clefts 1, 2 and 3. As expected, the Mackerel icefish and common carp proteins have increased solvent accessible surface area (Fig. 3c), which is consistent with the general trend (Fig. 2). However, the Mackerel icefish also has a very high level of protein flexibility especially in the cleft 2 region (Fig. 3d and Fig. S4b). The clustering analysis over each of the protein trajectories shows an increased population of conformations of LDHA away from the native structure for the Mackerel icefish (Fig. 3e), indicating a broader conformation space, especially at lower temperature (17 and 0 °C; Fig. 3e and Fig. S5a). This is in accord with the sequence difference that exists largely in cleft 2, where the common carp has one and Markerel icefish three more glycine than the Human protein (Fig. S5b). In summary, we infer an increased solvation for the LDHA protein of Antarctic fish by the occurrence of a larger number of polar amino acids on their surface; however, the protein also has enhanced internal dynamics, possibly a requirement for working in the extreme cold.

## Discussion

Understanding the evolution of proteins at the molecular level broadens our insight into the origin of species and the adaptation to their environment. The principle of natural evolution also gives insights into directed evolution in laboratory strategies for the optimization of functional enzymes and macromolecules to address challenges in health management and green chemistry. Instances of protein molecular adaptation have been recognized for several proteins, which have been studied across all three domains of life.^1–6,30–32^ Here we present a rather simple method to compare amino acid composition of two organisms at proteome, or rather sub-proteome level (since not all homologous proteins between two organisms have typically been identified yet). When we compare highly homologous proteins from hot- and cold-blooded model organisms, the method delivers a pattern of changes in the occurrence of amino acids which involves the balance of charge of the proteins, the number of polar residues on their surface and possibly also polypeptide chain dynamics. This is exciting because properties such internal protein dynamics and protein solubility have been recognized as important factors in protein thermal adaptation of individual proteins, comparing thermophilic with mesophilic species.^11–21^ For example, the decrease of polar residues and increase of hydrophobic residues with an increase of temperature is known to help enhance the stability of several proteins studied from thermophilic organisms.^33,34^ However, the thermal adaptation of proteins, especially enzymes and cell signaling proteins has been less consistent in terms of changes to their internal dynamics. There are cases where utilization of amino acids has clearly led to more flexible proteins at low temperature, but in other systems, the reverse or no difference is observed.^28–30, 35^

In this study, we investigated the conservation and shifts in the occurrence of amino acids in a pairwise comparison of protein sequences for sub-proteomes between human and chicken as representatives of hot-blooded and zebrafish and African claw frog as cold-blooded vertebrate organisms. The number of phenylalanine, histidine, glutamine, and to lesser extent cysteine differs little between the proteins of different organisms. Cysteine (for cell exterior proteins forming disulfide bonds) play a key role for protein stability, with Phe usually located in protein interiors, while His is commonly found at the surface regions, some located at protein-protein interactions sites. His and Cys, are also often highly conserved in chelating cofactors or ions. The conservation of these amino acids reflects on their role in maintaining the structural- and/or catalytic functions of the proteins. The exchange of aspartic acid with glutamic acid and exchange of lysine with arginine are both frequent. The observation that proteins usually tolerate/find advantages in substitutions that preserve amino acid similarity was made many decades ago.^e.g.36^ However, our analysis also indicates that cold-blooded organisms overall have a more negative net-charge per protein. While it is known that most proteins in bacteria have a lower isoelectric point, than human proteins, the reasons for these differences are not fully understood but may be related to issues with the long-term stability of polypeptide chains around acidic residues at higher temperatures.

Importantly, we noticed a distinction between proteins from warm-blooded and cold-blooded species considering the utilization of multiple polar-relative to non-polar amino acids. Specifically, we saw an increase in the occurrence of polar amino acids Ser, Asn and Thr as well as Ile, Met in fish and frog proteins; and a change in the occurrence of Leu, Ala, Gly, and Pro when comparing proteins from lower vertebrate species, such as fishes, with similar proteins from humans, other mammals and birds. The changes in occurrence are statistically significant with respect to the entire protein, but especially when seen relative to the number of amino acids of that type in a protein. We also note that these shifts in amino acid occurrence may in part arise by even functionally neutral changes of protein sequences, as the organisms compared differ considerably in their evolutionary distance from a common ancestor. However, we compared zebrafish and African clawed frog proteins to both human and chicken, which showed that while some of the changes are due to the evolution to higher level vertebrates, another part of the shift in amino acid utilization are associated with differences between proteins from cold and hot-blooded organisms. In general, the conservation as well as the shift in the occurrence of amino acid across these paired proteins reflect the complexity of the organism and the demands that its behavior and ultimately environment places on it. The latter comprises multiple factors, for example, the availability of food and nutrition, the metabolic level of an organism, and the difficulty of the biosynthesis of an amino acid as well as the pressure from nucleotide composition. Specifically, a correlation between the enrichment of hydrophobic amino acid groups, Ile, Val, Tyr, Leu and Trp upon an increase of optimal growth temperature, over a range of ~ 10 to 100 °C was noticed in comparing the proteomes of thermophilic bacteria by Berezovsky and colleagues.^33,34^ This team was also able to explain many of the trends with reference to nucleotide codon preferences which relate to the temperature and aerobic vs. anaerobic habitats of the Archaeal Bacterial genomes examined.^37^ Thus, we would not expect that a single overriding mechanism would explain all the shifts in amino acid occurrence.

Our results comparing hot- and cold-blooded vertebrate organisms suggest that the process of thermoadaptation may take into account the relationship between a protein’s structure, its internal dynamics and solubility. The amino acid composition of protein surfaces, linker and loops are known to play a critical role in determining these two features. Glycine, for example, has no side chain and is able to go through dihedral angles changes not allowed by the other residues. Mutations of other residues to glycine have been suggested to contribute to cold adaptation for enzymes such as Adenosine kinase and alkaline phosphatase, as these proteins require considerable dynamics for their catalytic cycles.^28,29^ At the same time, these changes could modulate protein association by changing a protein’s in protein internal dynamics and solvation. In the case of a change in dynamics, the reduced fluctuations of a protein at low body temperature, will already lessen the unfavorable entropy cost which is typically incurred when a protein binds a ligand or another protein. However, our sequence based predictions as well as molecular dynamics calculations suggest fish and frog proteins have considerably increased solvation, in accord with them having by having a greater number of polar amino acids, methionine and acidic amino acids. Importantly, we notice that our estimates for the change in the free energy of protein solvation tends to outweigh changes in protein internal dynamics, thus explaining why in the sub-proteome comparison Gly and Ala are changed to more bulky (less dynamic) polar residues, upon going from hot-to cold-blooded organisms. Still, the example of a specific protein the L-Lactate dehydrogenase A chain (LDHA) protein in Antarctic fish, which has both elevated protein dynamics and solvation, illustrates that similarly to some thermophilic proteins which are more dynamic than their mesophilic counterparts (noted above), there are exceptions. In this case, it is thought that the added flexibility is important for catalysis and occurred as an adaptation to the extreme cold of their ocean habitat.^28^

To end this paper, we would again emphasize that it is unlikely that one or a several few mechanisms can explain the changes in amino acid occurrence as identified in this study. More studies from a wider perspective of physicists, biologists and naturalists will be needed to give deeper insights into the underlying mechanisms. To “get the ball rolling”, at least at a vertebrate sub-proteome level, we propose that the change in amino acid occurrence in cold adaptation is at least in part related to the role of amino acids in determining protein internal dynamics as well as protein solvation, but that the effect of changes on protein solvation considerably outweighs the effects resulting from altered protein dynamics.

## Acknowledgements

This work is supported by a NIH R01 grant from the National Eye Institute R01EY029169 and previous grants from NIGMS (R01GM073071 and R01GM092851) to the Buck lab. The simulations were run at the Ohio Supercomputer Center (OSC), as well as local computing resource in the core facility for Advanced Research Computing at Case Western Reserve University.

## Author Contribution

Z.L. and M.B designed the studies, interpreted the data and wrote the manuscript. Z.L. performed the sequence analysis and computational modeling.

## Methods

### Comparative sequence analysis over sub-proteomes of human and model organisms

Protein sequences between Human (Homo sapiens) and multiple model vertebrate organisms, i.e. chimpanzee (Pan troglodytes), mouse (Mus musculus), chicken (Gallus gallus), African clawed frog (Xenopus laevis), and zebrafish (Danio rerio) were compared. All sequences of proteins of a species (including both “reviewed” and “un-reviewed” items) were downloaded from Uniprot database. Based on the Uniprot database, the total number of unique protein-coding genes of human is 20621 (gene counts). For chimpanzee, mouse, chicken, African clawed frog and zebrafish, the number is 23051, 21986, 18117, 43236, 25703 respectively (See Table S1). However, African clawed frog and zebrafish, have many more individual/species specific genes than mammals or chicken.^38^ With the downloaded sequences, we firstly analyzed the similar proteins between human and a model organism by matching the name of individual protein species. We identified 4136, 15674, 2681, 2016, 3383 proteins which occurred by name in both human and a model organism (chimpanzee/mouse/chicken/African clawed frog/zebrafish respectively). It should be realized that these numbers are not the real total number of shared proteins between human and a model organism. A considerable number of the protein sequences, in particular in the model organisms were not well reviewed on Uniprot websever and have not been linked to a specific protein species (marked as “uncharacterized protein”). Only the number of proteins paired between mouse and human should be close to the real number, as mouse has widely available reviewed protein sequences due to the extensive use of mice in the laboratory. Sequences of these selected proteins were pair-wise compared between the human and its homolog for each of the model organisms. To ensure a high degree of similarity we excluded protein pairs with a difference in sequence length >5%. As a result of this filter, we analyzed 3607, 14046, 1921, 1565, 2353 protein pairs between human and chimpanzee, mouse, chicken, African clawed frog, and zebrafish (see Table S1). However, we can regard the pairings as those between protein sequence homologs as the amino acid positional identity is greater than 30% for greater than 98% of the human-zebrafish and human-frog pairs, for example. For each protein, we compared the difference in absolute content for each one of the 20 amino acids (AA) in a protein pair (number of an amino acid in a protein scaled by the protein’s sequence length). Then the results from all other protein pairs are calculated similarly and collected in 9 different ranges (or bins) from −1% to 1% with an increment of 0.25%. Differences < −1% or >1% are added to the lowest or highest bin, respectively. The distribution of ΔC(AA) that arises is then over all the protein pairs (Fig. S1). The y—axis is the number of proteins in each bin for a given amino acid comparison, but could also be rescaled by dividing by the number of proteins, giving a % scale, making an easy comparison possible between amino acids and between the species comparisons. This % scale is the color bar plotted in Fig. 1c.

The content of an amino acid (ΔC(AA)) was compared one protein by another protein over the paired protein as selected above (for example, the 3607 paired proteins between human and chimpanzee). A two-tailed paired t-test was performed to check whether there is significant difference for the content of a specific amino acid between human and a model organism (p-value <0.05).

### Sequence-based energetic and entropy analysis

The average entropy of an amino acid in a polypeptide chain with the proteins sequence was obtained with values from Ref. 25; and the solvation free energy of an isolated amino acid side chain analogue was obtained from Ref. 26. The polypeptide chain entropy of a protein was estimated by adding up the entropy of the constituent amino acids. Proline was not included in the calculation of backbone entropy due to the unavailability of a value. The entropy was further averaged over the length of protein sequence (S_*bb*_ entropy per amino acid). For both entropy and solvation calculation, sequence lengths were adjusted by excluding the number of amino acids in each protein, for which no reference data are available (i.e., proline for entropy calculation; proline and charged residues for solvation calculation). The apparent entropy difference of a specific human and model organism protein pair was calculated as ΔS_*bb*_= S_*bb*_(model organism) – S_*bb*_(human). The solvation free energy of protein was estimated by adding up by the solvation free energy of the isolated amino acids. This estimates the upper limit of a protein solvation free energy. The difference in protein solvation free energy between a protein pair was calculated as ΔG_*sol*_= G_*sol*_(model organism) – G_*sol*_(human). Proline and Charged residues were not included in the calculation of solvation free energy due to the unavailability of experimental data for these amino acids. The average of of ΔSave = {ΔS_1_, ΔS_2_, …ΔS_N_}/N and ΔGave ={ΔG_1_, ΔG_2_, … ΔG_N_}/N of N pair-wise proteins between human and model organisms, as well their standard variation of are in plotted Fig. 2 and Fig. S3.

### Molecular modeling of 166 paired proteins between human and zebrafish

We analyzed 2353 human proteins and as many of their homologs we could find in zebrafish. Sequence alignment between zebrafish and human proteins were carried out with bioPython with the Clustalw module. Of 2353 proteins, 644 have available PDB structures for the human proteins. Selectively, we chose structures that have a resolution better than 5 Å, cover at least 80% protein sequence, and do not have extensive flexible regions such as long loops **(> N residues?)**. This reduced the set to 166 soluble proteins. On average the protein pairs had high sequence identity and thus expected structural homology (106 of 166 pairs with >70% identity; all but 10 >50% identity; the remaining 10 had >36% sequence identity). A full list of proteins studied is given in Table S4. Modeller was used to build the initial structure of each protein, adding missing residues and short loops.^39^ For human proteins, N-terminal and C-terminal residues were added automatically if the missing segments are less than 5 residues long in a PDB. If there were more than 5 residues missing, the N- or the C-terminal residue of a protein was kept the same as in a PDB structure. The structure of the pair-wise zebrafish protein was constructed based on the corresponding human protein PDB with Modeller. The modeled protein systems were subjected to standard molecular dynamics simulations each of 100 ns (see details below). Disulfide bonds were patched with module DISU in the CHARMM force field as needed for specific proteins. Even though the simulations are short, the great majority of simulations should equilibrate since proteins with long loops were excluded. All these simulations were carried out at 310K. The results were analyzed by comparing the properties of the protein pairs in a manner similar to the sequence-based energetic analysis. Specifically, ΔRMSF (Root mean squared fluctuation), ΔE (Poisson Boltzmann electrostatic interaction), ΔSASA (Solvent available surface area), and ΔH-bonds (number of hydrogen bonds between solvent and protein) were calculated and the latter three parameters were scaled by the length of the protein, i.e. are given per residue. All are shown for the 166 simulated proteins with a boxplot in Fig. 2c.

### Case study of L-Lactate dehydrogenase A chain (LDHA)

In the sequence analysis of LDHA, 17 Antarctic fishes were considered for nothothenioidei (Antarctic spiny plunderfish, Patagonian toothfish, Antarctic cod, Humped rockcod, Charcot’s dragonfish, Yellowbelly rockcod, Gaudy notothen, Mackerel icefish, Emerald rockcod, Blackfin icefish, Ocellated icefish, Rockcod, Bald rockcod, Patagonian blennie, Maori cod, Black southern cod, Antarctic eelpout); 10 fishes (excluding the Antarctic fish) were used for actinopterygii, fish living in fresh or more moderate temperature salt water (Pacific barracuda, Pelican barracuda, Lucas barracuda, Zebrafish, Killifish, Common carp, Long-jawed mudsucker, Shortjaw mudsucker, Blackeye goby, Spiny dogfish); Apart from the fish, 8 species were used for cold-blooded organisms (African claweded frog, Axolotl, Eastern fence lizard, Ball python, Florida scrub lizard, Chinese soft-shelled turtle, Red-eared slider turtle, American alligator) and 10 species for mammals (Gray short-tailed opossum, Rabbit, Pig, Wild yak, Bovine, Crab-eating macaque, Sumatran orangutan, Chimpanzee, Mouse, Rat, Human) were used as hot-blooded organisms for the amino acid occurrence analysis. Our standard simulation procedure was applied to the LDHA protein. PDB entry 4ajp, 1v6a and 2v65 were used for human, common carp and Mackerel icefish protein respectively. Each simulation system was performed in triplicate for 500 ns at three different temperatures - 310K (37°C), 290K (17°C) and 273K (0°C),

### General simulation parameters for all modeling

All proteins were solvated by TIP3P water with 150 mM NaCl; all simulations were performed with the NAMD/2.12 package.^40^ For simulations of 100 ns in length, they were performed on the GPU cluster at Case Western Reverse University or at the Ohio supercomputer center (OSC). A time step of 2 fs was employed. The SHAKE algorithm was applied for all covalent bonds to hydrogen. A Langevin thermostat of 310 K and a semi-isotropic Langevin scheme at 1 bar was used. The CHAMRM36m force field was used in all simulations.^41^ The van der Waals (vdW) potential was cut off at 12 Å and smoothly shifted to zero between 10 and 12 Å. The long-range electrostatic interactions were calculated with the Particle-Mesh Ewald (PME) method.

