## supporting_information for "A Proteome-scale Analysis of Vertebrate Protein Amino Acid Occurrence: Thermoadaptation shows a Correlation with Protein Solvation but less so with Dynamics"

Zhen-lu Li [1*] and Matthias Buck [1,2*]

[1]Department of Physiology and Biophysics, Case Western Reserve University, School of Medicine, 10900 Euclid Avenue, Cleveland, Ohio 44106, U. S. A. [2]Department of Pharmacology; Department of Neurosciences, Case Western Reserve University, School of Medicine, 10900 Euclid Avenue, Cleveland, Ohio 44106, U. S. A.

Table S1: Information for proteomes of human, chimpanzee, mouse, chicken, African clawed frog and zebrafish. The total number of proteins of an organism which are homologous to human proteins is unavailable/remains to be determined. Based on the current database of proteins by species in the Uniprot websever (end July 2020), we identify corresponding homologous proteins between human and a model organism (column 7). For analysis, homologous proteins are considered only if the sequence length difference <5% between human protein and its homolog (column 8). As noted these two criteria ensure that > 98% of the protein are > 30% sequence identical and can thus be considered homologous.

| Organism | Total Gene Count | Individual -Specific Proteins | Total Protein Counts | Reviewed Items | Total homologous proteins to human | Homologous Proteins identified in the study | Homologous Proteins (<5% sequence difference) |
|---|---|---|---|---|---|---|---|
| Human | 20621 | Low | 75004 | 20353 | N/A | | |
| Chimpanzee | 23051 | Low | 134654 | 693 | Unknown | 4136 | 3607 |
| Mouse | 21986 | Low | 86532 | 17042 | Unknown | 15674 | 14046 |
| Chicken | 18117 | Low | 34748 | 2295 | Unknown | 2681 | 1921 |
| African clawed frog | 43236 | High | 57801 | 3451 | Unknown | 2016 | 1565 |
| zebrafish | 25703 | High | 46847 | 3455 | Unknown | 3383 | 2353 |

Table S2a: Stastitical analysis of average content of an amino acid for paired protein between an organism and human (row 1 and row 2 for each comparison). The absolute content of an amino acid are given in % in the table. The difference and significance (p-value) are given in row 3, 4.

| | Organism | A | C | D | E | F | G | H | I | K | L | M | N | P | Q | R | S | T | V | W | Y |
|---|---|---|---|---|---|---|---|---|---|---|---|---|---|---|---|---|---|---|---|---|---|
| N=3607 | Chimpanzee | 6.95 +/- 2.74 | 2.86 +/- 2.94 | 4.43 +/- 1.82 | 6.68 +/- 2.69 | 3.94 +/- 1.7 | 6.69 +/- 3.0 | 2.79 +/- 1.64 | 4.36 +/- 2.05 | 5.73 +/- 2.88 | 10.02+/- 3.28 | 2.21 +/- 1.12 | 3.52 +/- 1.58 | 6.00 +/- 3.15 | 4.52 +/- 1.89 | 5.65 +/- 2.27 | 7.88 +/- 2.54 | 5.26 +/- 1.66 | 5.98 +/- 2.02 | 1.30 +/- 0.91 | 2.92 +/- 1.76 |
| | Human | 6.95 +/- 2.75 | 2.85 +/- 2.93 | 4.44 +/- 1.84 | 6.70 +/- 2.76 | 3.94 +/- 1.72 | 6.68 +/- 3.0 | 2.78 +/- 1.64 | 4.37 +/- 2.05 | 5.74 +/- 2.88 | 10.01+/- 3.29 | 2.22 +/- 1.12 | 3.53 +/- 1.58 | 5.99 +/- 3.16 | 4.52 +/- 1.88 | 5.63 +/- 2.27 | 7.87 +/- 2.55 | 5.26 +/- 1.66 | 5.98 +/- 2.03 | 1.29 +/- 0.92 | 2.92 +/- 1.76 |
| | Difference | 0 | 0.01 | -0.01 | -0.02 | 0 | 0.01 | 0.01 | -0.01 | -0.01 | 0.01 | -0.01 | -0.01 | 0.01 | 0 | 0.02 | 0.01 | 0 | 0 | 0.01 | 0 |
| | p-value | 0.2059 | 0.0094 | 0.0005 | 0.0015 | 0.5546 | 0.204 | 0.0052 | 0.0486 | 0.0731 | 0.1735 | 0.5402 | 0.4971 | 0.0037 | 0.7655 | 0.0029 | 0.7083 | 0.9671 | 0.8242 | 0.1268 | 0.1808 |
| N=14046 | mouse | 7.19 +/- 2.4 | 2.27 +/- 1.81 | 4.8 +/- 1.64 | 6.83 +/- 2.65 | 3.82 +/- 1.62 | 6.6 +/- 2.56 | 2.51 +/- 1.16 | 4.33 +/- 1.83 | 5.71 +/- 2.62 | 10.14+/- 2.83 | 2.29 +/- 1.03 | 3.49 +/- 1.41 | 5.96 +/- 2.74 | 4.55 +/- 1.76 | 5.77 +/- 2.15 | 7.93 +/- 2.47 | 5.18 +/- 1.55 | 6.21 +/- 1.83 | 1.28 +/- 0.86 | 2.81 +/- 1.33 |
| | Human | 7.3 +/- 2.61 | 2.25 +/- 1.78 | 4.76 +/- 1.63 | 6.87 +/- 2.69 | 3.82 +/- 1.63 | 6.7 +/- 2.64 | 2.48 +/- 1.16 | 4.41 +/- 1.91 | 5.74 +/- 2.71 | 10.14+/- 2.85 | 2.27 +/- 1.03 | 3.55 +/- 1.5 | 6.03 +/- 2.86 | 4.52 +/- 1.76 | 5.77 +/- 2.21 | 7.75 +/- 2.45 | 5.09 +/- 1.57 | 6.13 +/- 1.84 | 1.28 +/- 0.86 | 2.82 +/- 1.33 |
| | Difference | -0.11 | 0.02 | 0.04 | -0.04 | 0 | -0.1 | 0.03 | -0.08 | -0.03 | 0 | 0.02 | -0.06 | -0.07 | 0.03 | 0 | 0.18 | 0.09 | 0.08 | 0 | -0.01 |
| | p-value | <0.0001 | <0.0001 | <0.0001 | <0.0001 | 0.7717 | <0.0001 | <0.0001 | <0.0001 | <0.0001 | 0.2751 | <0.0001 | <0.0001 | <0.0001 | <0.0001 | 0.5253 | <0.0001 | <0.0001 | <0.0001 | 0.5213 | 0.0001 |
| N=1921 | chicken | 7.15 +/- 2.49 | 2.13 +/- 1.49 | 5.04 +/- 1.8 | 7.1 +/- 2.65 | 3.96 +/- 1.68 | 6.32 +/- 2.55 | 2.42 +/- 1.16 | 4.91 +/- 1.78 | 6.37 +/- 2.76 | 9.69 +/- 2.77 | 2.49 +/- 1.02 | 3.85 +/- 1.39 | 5.33 +/- 2.43 | 4.35 +/- 1.7 | 5.49 +/- 2.08 | 7.41 +/- 2.51 | 5.10 +/- 1.50 | 6.31 +/- 1.91 | 1.22 +/- 0.84 | 3.06 +/- 1.32 |
| | Human | 7.07 +/- 2.44 | 2.08 +/- 1.48 | 5.03 +/- 1.75 | 7.04 +/- 2.68 | 3.98 +/- 1.68 | 6.42 +/- 2.51 | 2.44 +/- 1.16 | 4.79 +/- 1.83 | 6.22 +/- 2.76 | 9.95 +/- 2.83 | 2.45 +/- 1.05 | 3.76 +/- 1.42 | 5.52 +/- 2.53 | 4.44 +/- 1.7 | 5.48 +/- 2.02 | 7.39 +/- 2.39 | 5.14 +/- 1.49 | 6.29 +/- 1.83 | 1.24 +/- 0.86 | 2.98 +/- 1.29 |
| | Difference | 0.08 | 0.05 | 0.01 | 0.06 | -0.02 | -0.1 | -0.02 | 0.12 | 0.15 | -0.26 | 0.04 | 0.09 | -0.19 | -0.09 | 0.01 | 0.02 | -0.04 | 0.02 | -0.02 | 0.08 |
| | p-value | 0.0555 | <0.0001 | 0.7499 | 0.0047 | 0.1016 | 0.0002 | 0.1955 | <0.0001 | <0.0001 | <0.0001 | 0.024 | <0.0001 | <0.0001 | <0.0001 | 0.675 | 0.4897 | 0.142 | 0.4642 | 0.0032 | <0.0001 |
| N=1565 | African Clawed Frog | 6.34 +/- 1.87 | 2.18 +/- 1.52 | 5.22 +/- 1.78 | 7.01 +/- 2.69 | 3.96 +/- 1.65 | 5.93 +/- 2.26 | 2.55 +/- 1.17 | 5.17 +/- 1.64 | 6.56 +/- 2.6 | 9.77 +/- 2.55 | 2.57 +/- 1.06 | 4.12 +/- 1.32 | 4.92 +/- 2.06 | 4.53 +/- 1.7 | 5.22 +/- 1.86 | 7.9 +/- 2.41 | 5.21 +/- 1.37 | 6.21 +/- 1.75 | 1.20 +/- 0.80 | 3.11 +/- 1.31 |
| | Human | 7.16 +/- 2.34 | 2.12 +/- 1.5 | 5.03 +/- 1.77 | 7.05 +/- 2.75 | 3.91 +/- 1.62 | 6.36 +/- 2.37 | 2.51 +/- 1.14 | 4.65 +/- 1.78 | 6.15 +/- 2.7 | 10.19+/- 2.71 | 2.41 +/- 1.05 | 3.64 +/- 1.41 | 5.45 +/- 2.46 | 4.51 +/- 1.71 | 5.61 +/- 2.0 | 7.46 +/- 2.33 | 5.00 +/- 1.40 | 6.29 +/- 1.82 | 1.24 +/- 0.82 | 2.96 +/- 1.3 |
| | Difference | -0.82 | 0.06 | 0.19 | -0.04 | 0.05 | -0.43 | 0.04 | 0.52 | 0.41 | -0.42 | 0.16 | 0.48 | -0.53 | 0.02 | -0.39 | 0.44 | 0.21 | -0.08 | -0.04 | 0.15 |
| | p-value | <0.0001 | <0.0001 | <0.0001 | 0.0724 | 0.0022 | <0.0001 | 0.0262 | <0.0001 | <0.0001 | <0.0001 | <0.0001 | <0.0001 | <0.0001 | 0.4124 | <0.0001 | <0.0001 | <0.0001 | 0.0097 | <0.0001 | <0.0001 |
| N=2353 | Zebra-fish | 6.67 +/- 2.15 | 2.15 +/- 1.54 | 5.23 +/- 1.87 | 6.77 +/- 2.63 | 4.03 +/- 1.71 | 6.23 +/- 2.52 | 2.53 +/- 1.2 | 4.80 +/- 1.74 | 6.13 +/- 2.72 | 9.99 +/- 2.78 | 2.70 +/- 1.19 | 3.81 +/- 1.33 | 4.90 +/- 2.15 | 4.50 +/- 1.85 | 5.48 +/- 2.06 | 7.72 +/- 2.47 | 5.27 +/- 1.67 | 6.53 +/- 1.9 | 1.23 +/- 0.85 | 2.94 +/- 1.34 |
| | Human | 7.19 +/- 2.46 | 2.09 +/- 1.52 | 4.91 +/- 1.81 | 6.86 +/- 2.74 | 4.01 +/- 1.7 | 6.52 +/- 2.54 | 2.49 +/- 1.13 | 4.67 +/- 1.87 | 6.04 +/- 2.86 | 10.49+/- 2.91 | 2.46 +/- 1.17 | 3.61 +/- 1.47 | 5.31 +/- 2.25 | 4.50 +/- 1.77 | 5.62 +/- 2.12 | 7.23 +/- 2.33 | 5.03 +/- 1.66 | 6.34 +/- 1.85 | 1.29 +/- 0.88 | 2.96 +/- 1.35 |
| | Difference | -0.52 | 0.06 | 0.32 | -0.09 | 0.02 | -0.29 | 0.04 | 0.13 | 0.09 | -0.5 | 0.24 | 0.2 | -0.41 | 0 | -0.14 | 0.49 | 0.24 | 0.19 | -0.06 | -0.02 |
| | p-value | <0.0001 | <0.0001 | <0.0001 | 0.0003 | 0.2631 | <0.0001 | 0.0117 | 0.0001 | 0.0063 | <0.0001 | <0.0001 | <0.0001 | <0.0001 | 0.7923 | <0.0001 | <0.0001 | <0.0001 | <0.0001 | <0.0001 | 0.1887 |

Table S2b: Same as table S1a but for zebrafish and African clawed frog compared to Chicken.

| | Organism | A | C | D | E | F | G | H | I | K | L | M | N | P | Q | R | S | T | V | W | Y |
|---|---|---|---|---|---|---|---|---|---|---|---|---|---|---|---|---|---|---|---|---|---|
| N=1036 | African clawedC Frog | 6.50 +/- 2.06 | 2.02 +/- 1.30 | 5.19 +/- 1.68 | 7.05 +/- 2.55 | 3.94 +/- 1.62 | 6.00 +/- 2.34 | 2.52 +/- 1.18 | 5.22 +/- 1.68 | 6.77 +/- 2.64 | 9.64 +/- 2.59 | 2.67 +/- 1.07 | 4.04 +/- 1.31 | 4.90 +/- 2.05 | 4.46 +/- 1.71 | 5.32 +/- 2.10 | 7.56 +/- 2.46 | 5.23 +/- 1.42 | 6.35 +/- 1.85 | 1.22 +/- 0.88 | 3.09 +/- 1.76 |
| | Chicken | 7.32 +/- 2.43 | 1.98 +/- 1.3 | 5.00 +/- 1.70 | 7.18 +/- 2.58 | 3.89 +/- 1.61 | 6.25 +/- 2.4 | 2.46 +/- 1.18 | 4.89 +/- 1.71 | 6.55 +/- 2.68 | 9.79 +/- 2.73 | 2.54 +/- 1.02 | 3.72 +/- 1.35 | 5.14 +/- 2.3 | 4.37 +/- 1.72 | 5.69 +/- 2.25 | 7.24 +/- 2.42 | 5.02 +/- 1.56 | 6.38 +/- 1.95 | 1.24 +/- 0.87 | 3.02 +/- 1.26 |
| | Difference | -0.82 | 0.04 | 0.19 | -0.13 | 0.05 | -0.25 | 0.06 | 0.33 | 0.22 | -0.15 | 0.13 | 0.32 | -0.24 | 0.09 | -0.37 | 0.32 | 0.21 | -0.03 | -0.02 | 0.07 |
| | p-value | <0.0001 | 0.0068 | <0.0001 | 0.0002 | 0.0335 | <0.0001 | 0.0008 | <0.0001 | <0.0001 | <0.0001 | <0.0001 | <0.0001 | <0.0001 | 0.0014 | <0.0001 | <0.0001 | <0.0001 | 0.31116 | 0.0172 | 0.0003 |
| N=1499 | Zebra-fish | 6.73 +/- 2.13 | 2.08 +/- 1.40 | 5.28 +/- 1.69 | 6.91 +/- 2.51 | 3.99 +/- 1.69 | 6.04 +/- 2.16 | 2.56 +/- 1.12 | 4.81 +/- 1.61 | 6.19 +/- 2.58 | 9.80 +/- 2.64 | 2.64 +/- 1.04 | 3.84 +/- 1.27 | 4.97 +/- 1.97 | 4.52 +/- 1.72 | 5.52 +/- 1.89 | 7.78 +/- 2.40 | 5.34 +/- 1.41 | 6.61 +/- 1.83 | 1.19 +/- 0.84 | 2.91 +/- 1.26 |
| | Chicken | 7.24 +/- 2.39 | 2.07 +/- 1.37 | 4.98 +/- 1.66 | 7.12 +/- 2.59 | 3.96 +/- 1.69 | 6.24 +/- 2.26 | 2.48 +/- 1.10 | 4.93 +/- 1.72 | 6.33 +/- 2.61 | 9.97 +/- 2.66 | 2.47 +/- 1.02 | 3.72 +/- 1.34 | 5.19 +/- 2.28 | 4.40 +/- 1.64 | 5.62 +/- 2.09 | 7.27 +/- 2.33 | 5.04 +/- 1.48 | 6.44 +/- 1.84 | 1.23 +/- 0.86 | 3.0+/- 1.29 |
| | Difference | -0.51 | 0.01 | 0.30 | -0.21 | 0.03 | -0.20 | 0.08 | -0.12 | -0.14 | -0.17 | 0.17 | 0.12 | -0.22 | 0.12 | -0.01 | 0.51 | 0.30 | 0.17 | -0.04 | -0.09 |
| | p-value | <0.0001 | 0.4972 | <0.0001 | <0.0001 | 0.2619 | <0.0001 | 0.0001 | 0.0001 | 0.0003 | <0.0001 | <0.0001 | <0.0001 | <0.0001 | <0.0001 | 0.0048 | <0.0001 | <0.0001 | <0.0001 | <0.0001 | <0.0001 |

Figure S1a: Conservation and shift in amino acid content of proteins, comparing model organisms and human sub-proteomes. Distribution of ΔC over 9 bins from -1% to 1% with an increment of 0.25% (Similar to Fig. 1b). Listed for 20 amino acids. Comparison is done between (a) Chimpanzee and human; (b) mouse and human; (c) chicken and human; (d) African clway frog and human; (e) zebrafish and human.

a

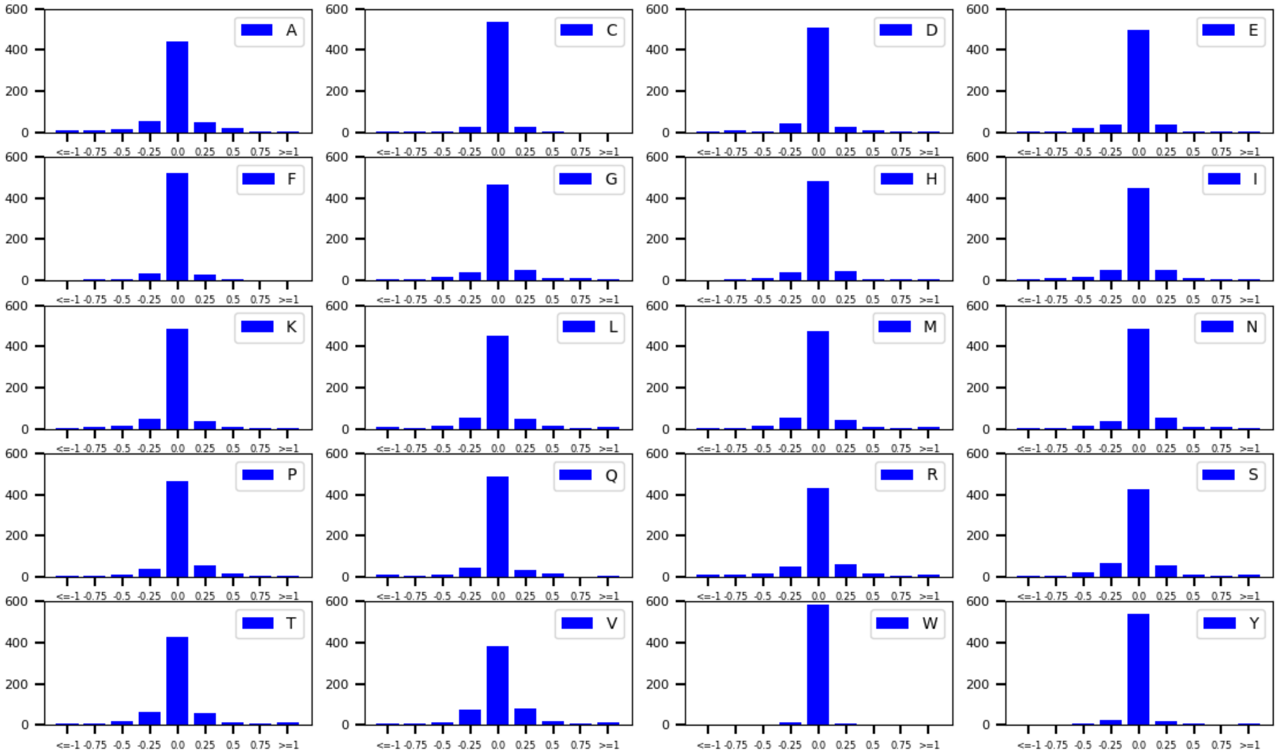

b

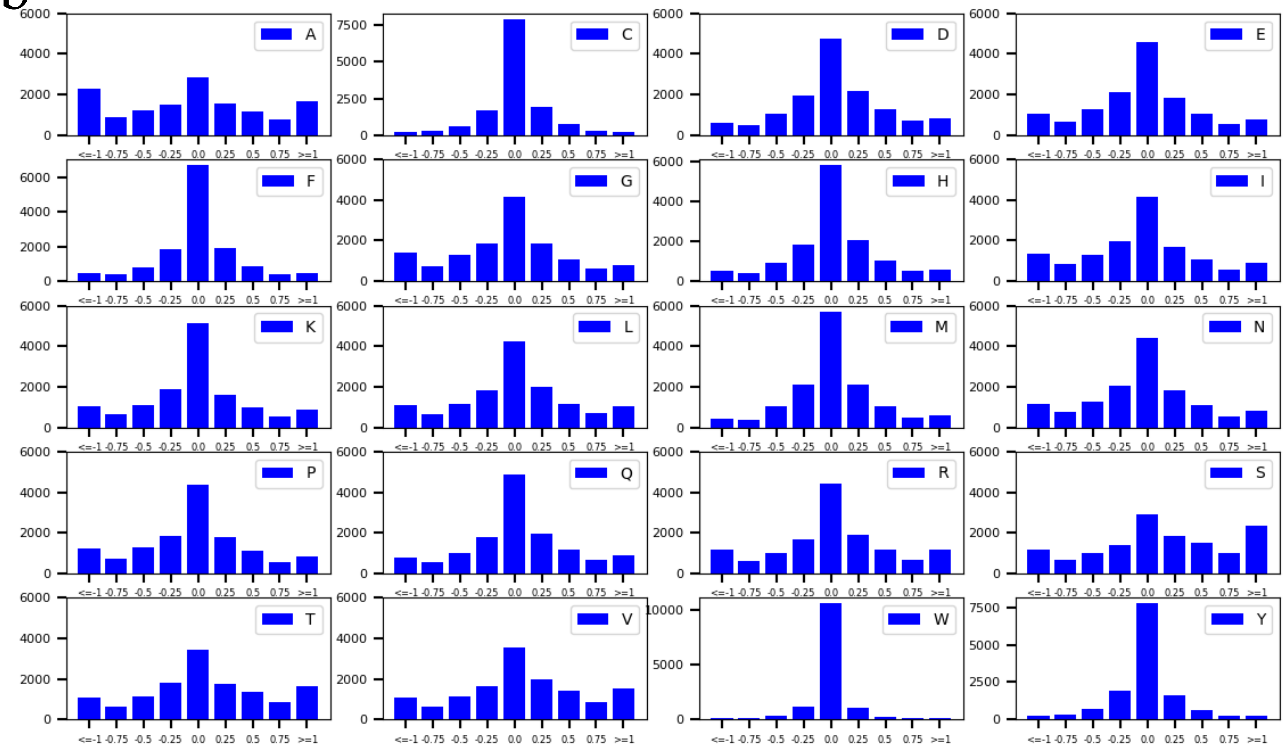

c

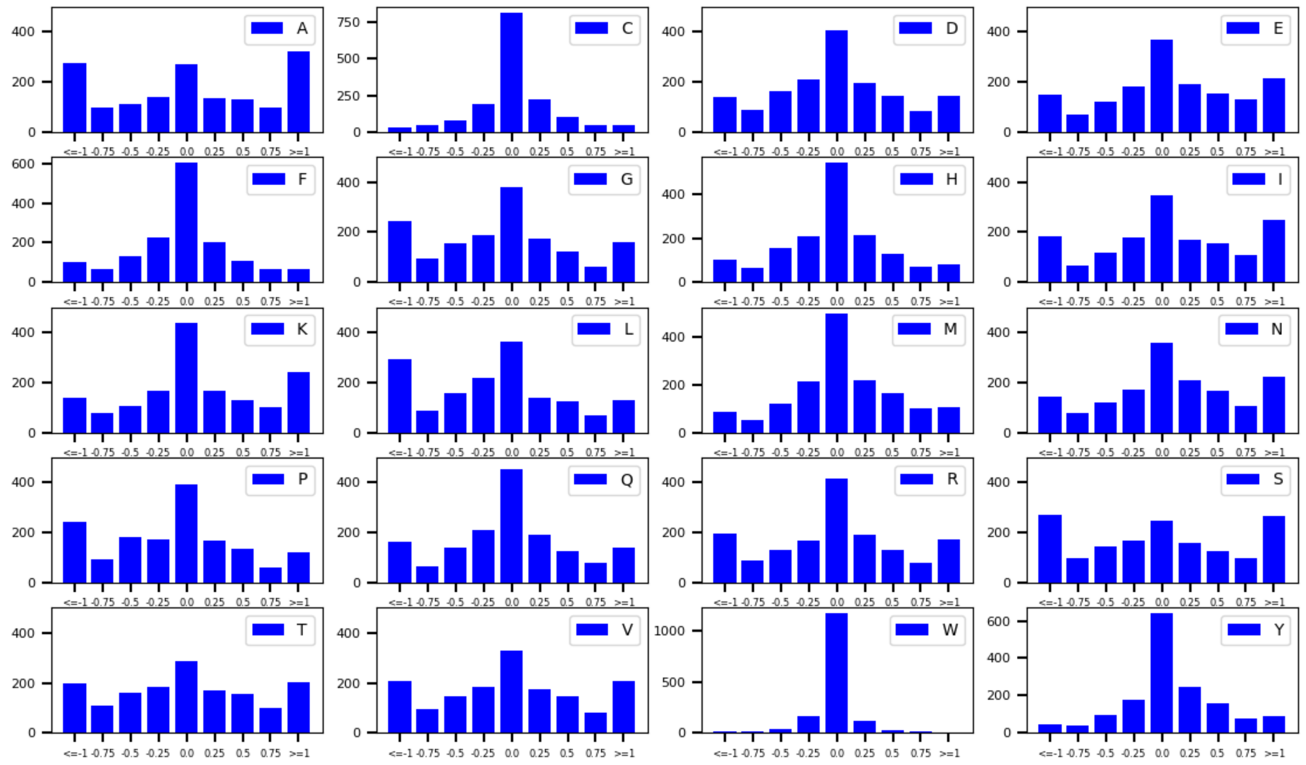
A
C
D
E
F
G
H
I
K
L
M
N
P
Q
R
S
T
V
W
Y
<=-1 -0.75 -0.5 -0.25 0.0 0.25 0.5 0.75 >=1

d

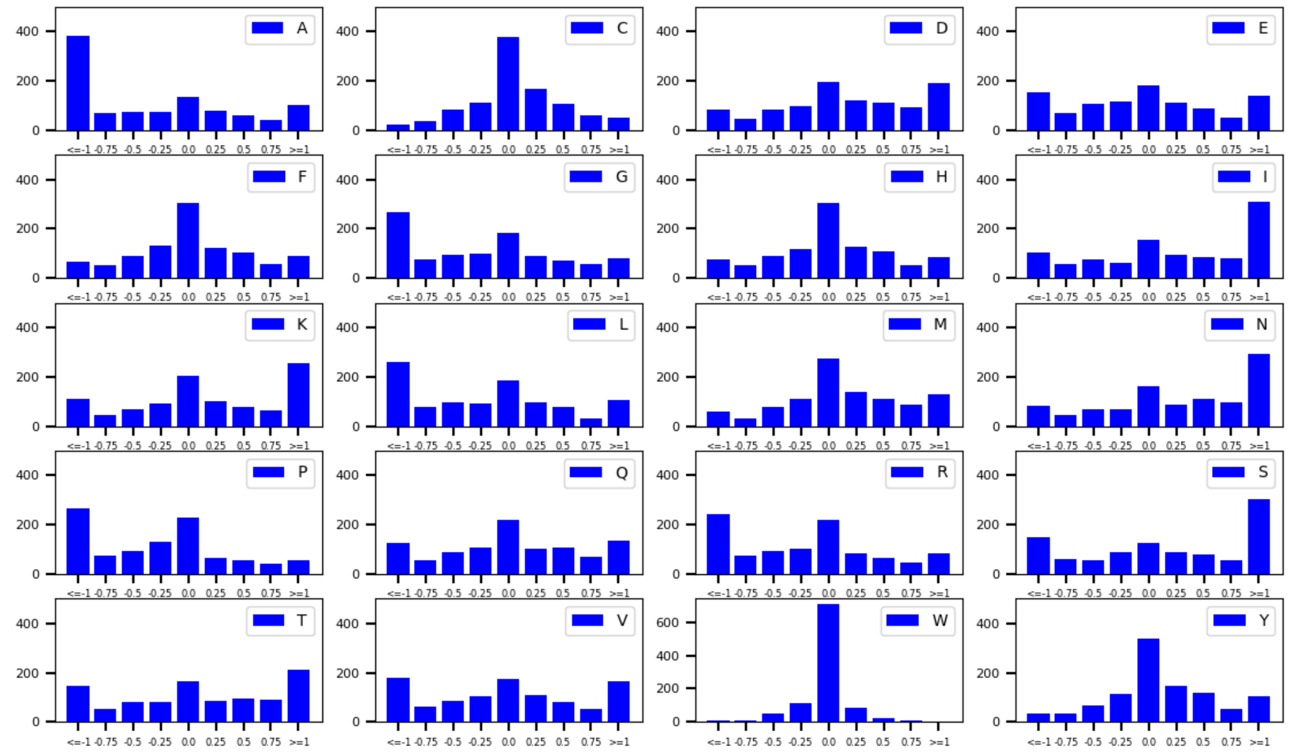
A
C
D
E
F
G
H
I
K
L
M
N
P
Q
R
S
T
V
W
Y
<=-1 -0.75 -0.5 -0.25 0.0 0.25 0.5 0.75 >=1

e

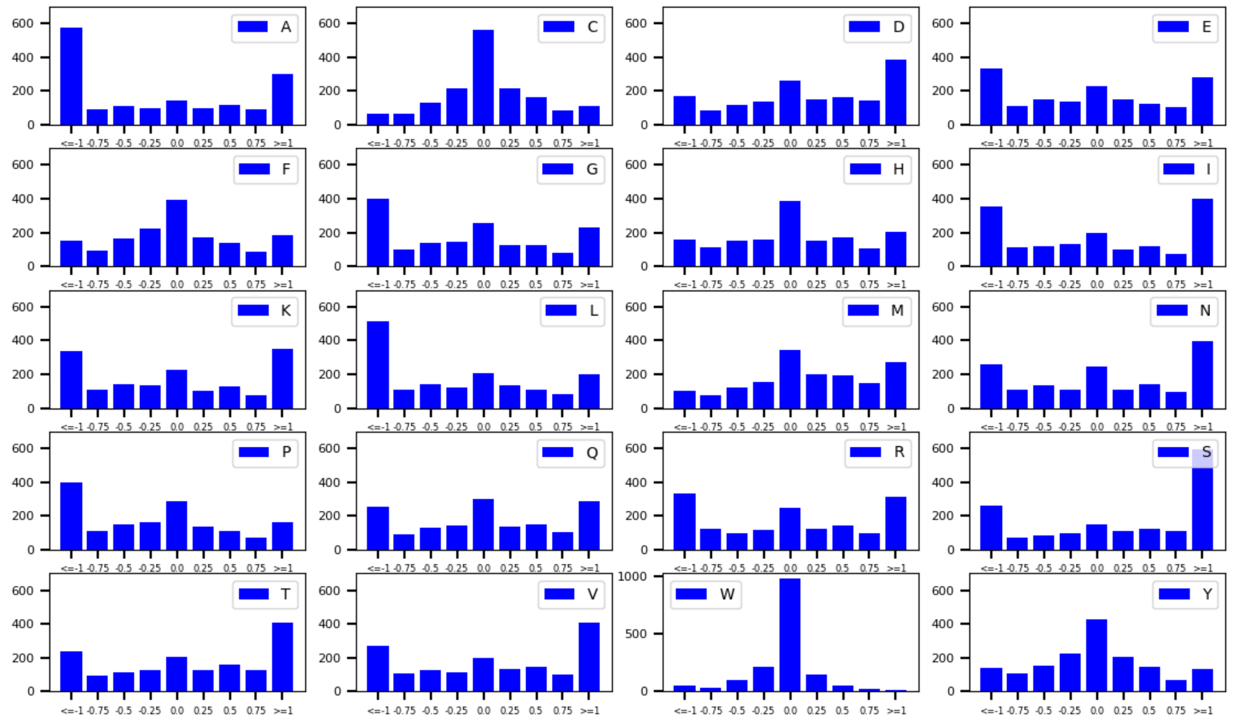

Figure S1b: Same as Fig S1a but for zebrafish and African clawed frog compared to chicken. Comparison is done between (a) African clawed frog and chicken; (b) zebrafish and chicken.

a

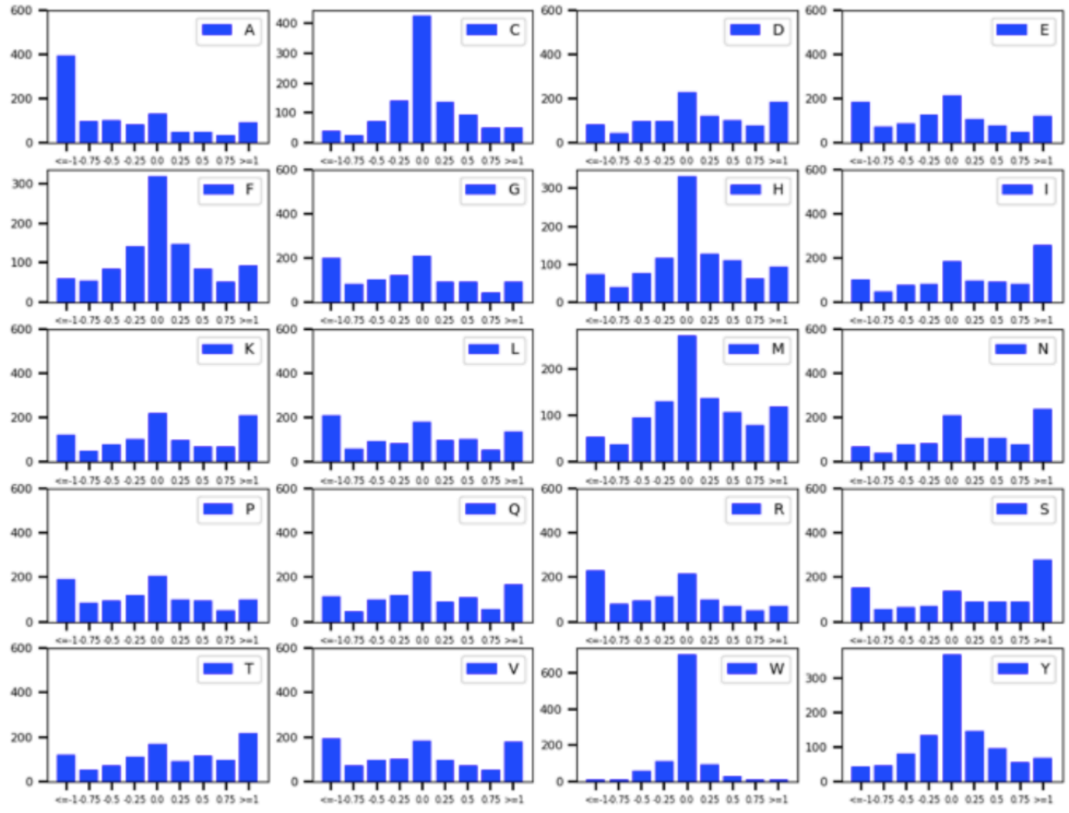

b

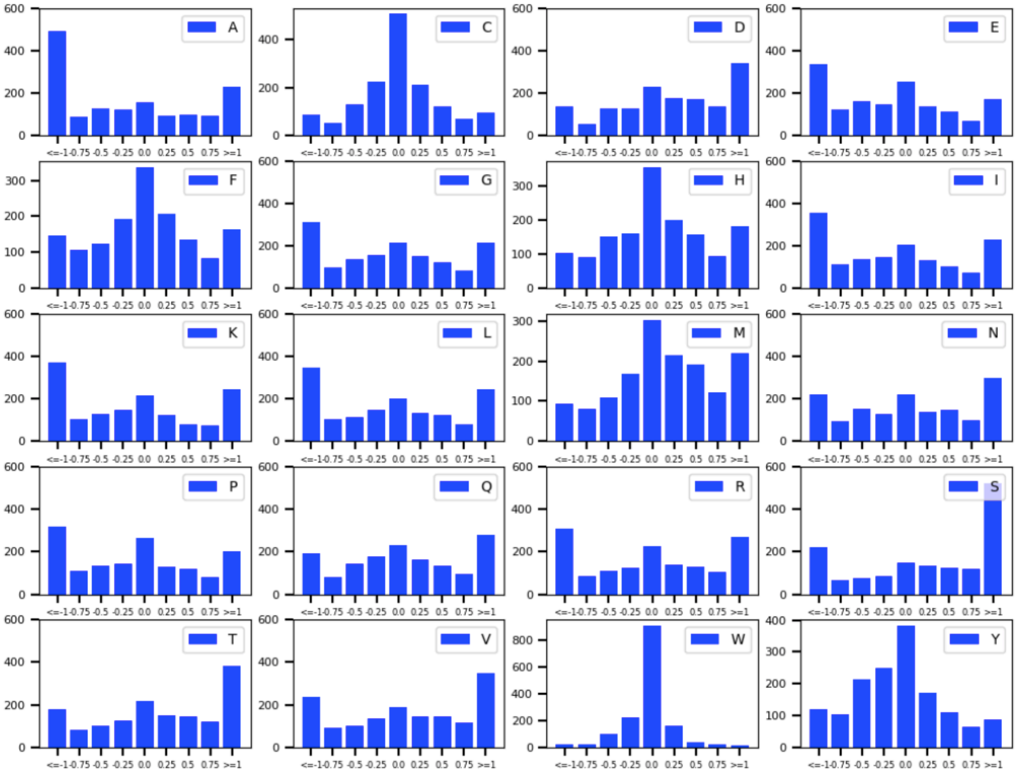

Table S3: Statistical analysis of contents of amino acids grouped into 5 types between chicken and African clawed frog or zebrafish. 1036 and 1499 paired proteins are compared for the analysis. The difference in absolute content and the significance (p-value) are added under each column.

| | Organism | G | A/I/L | K/R | D/E | N/S/T/M |
|---|---|---|---|---|---|---|
| N=1036 | African Clawed Frog | 6.00% | 21.36% | 12.08% | 12.23% | 19.81% |
| | chicken | 6.25% | 22.01% | 12.24% | 12.08% | 18.51% |
| | Difference Δ | **-0.25%** | **-0.65%** | **-0.16%** | **0.06%** | **1.0%** |
| | p-value | <0.0001 | <0.0001 | <0.0001 | 0.0753 | <0.0001 |
| N=1499 | zebrafish | 6.04% | 21.33% | 11.72% | 12.19% | 19.6% |
| | chicken | 6.24% | 22.14% | 11.95% | 12.10% | 18.49% |
| | Difference Δ | **-0.2%** | **-0.81%** | **-0.23%** | **0.09%** | **1.11%** |
| | p-value | <0.0001 | <0.0001 | <0.0001 | 0.0029 | <0.0001 |

Figure S2. Difference (in %) of (a) basic and (b) acidic amino acid content between homolog protein pairs in a model organism and the human protein. The distribution is over 3607, 14046, 1921, 1565, and 2353 proteins for chimapanzee, mouse, chicken, Afrian claw frog, and zebrafish respectively. Supposing a charge of +1 for lysine and arginine and -1 for aspartic and glutamic acid, on average the African clawed frog has a reduced positive charge by -0.14 units per protein and more negative charge by -0.43 per protein in comparison to the corresponding human protein. For zebrafish, the values are -0.28 (less basic amino acids) and -0.66 (more acidic amino acids). In contrast to the observed difference of -0.57 and -0.94 in net charge per protein for African clawed frog and zebrafish, the difference in average net charge between chimpanzee, mouse, chicken and human is +0.16, -0.11 and +0.32 charge units per protein respectively.

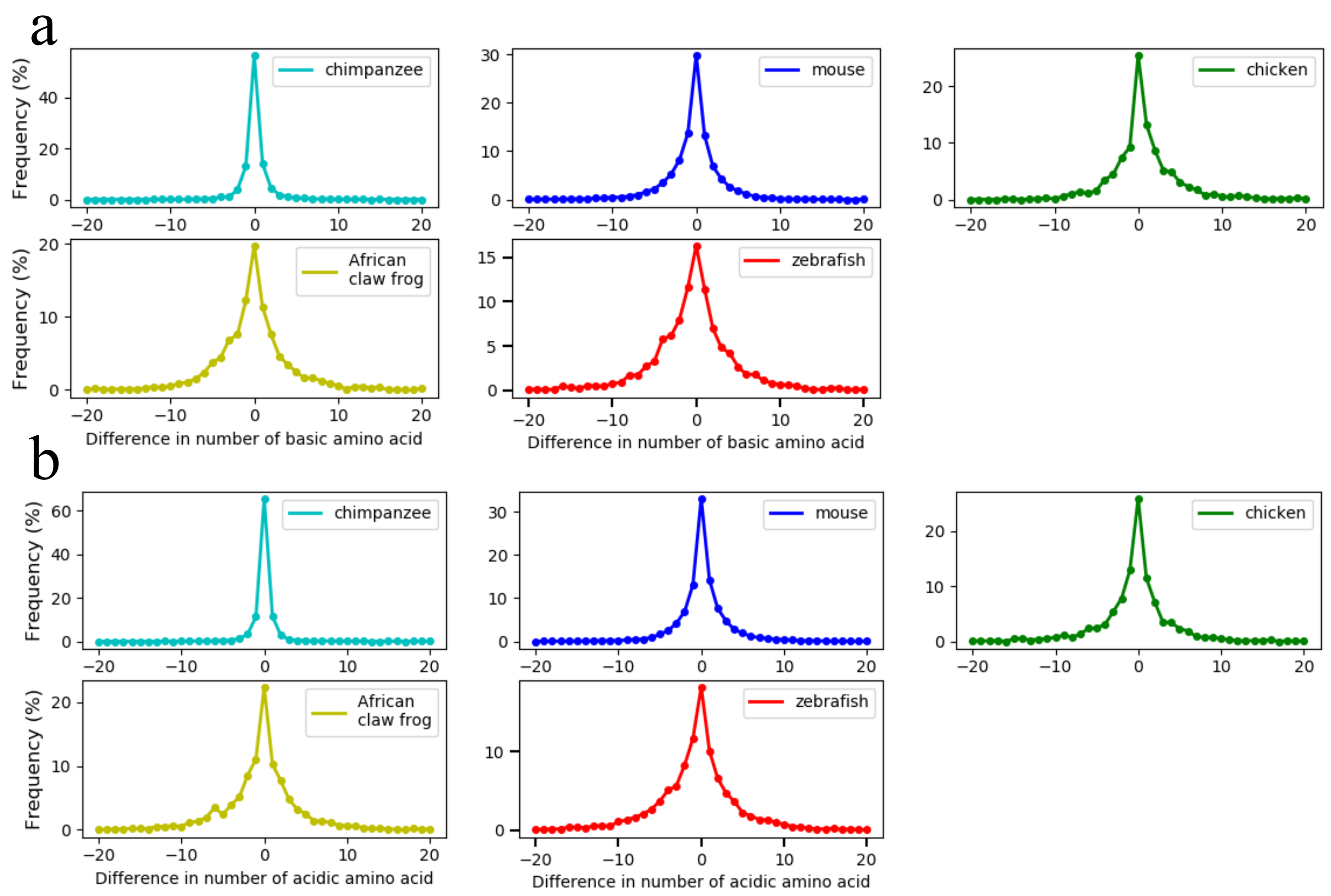

Figure S3: Sequence based backbone entropy and solvation free energy analaysis for homologous proteins between model organism and human proteins. (a) The table list the values of available experimental value of amino acid backbone entropy (*) and solvation free energy (**) from ref. 39, 40. (b) Entropy and (c) enthalpy difference over 3607, 14046, 1921, 1565, and 2353 proteins between chimapanzee, mouse, chicken, Afrian clawed frog, and zebrafish respectively (see methods for details).

a

| | $\Delta S_{bb}$(cal/K/mol)* | $\Delta G_{sol}$(kcal/mol)** | | $\Delta S_{bb}$(cal/K/mol)* | $\Delta G_{sol}$(kcal/mol)** |
|---|---|---|---|---|---|
| ALA | 4.1 | 1.94 | TYR | 3.4 | -6.11 |
| VAL | 2.18 | 1.99 | PHE | 3.4 | -0.76 |
| LEU | 3.4 | 2.28 | TRP | 3.4 | -5.88 |
| ILE | 2.18 | 2.15 | HIS | 3.4 | -10.27 |
| SER | 3.4 | -5.06 | GLY | 6.5 | - |
| THR | 3.4 | -4.88 | ARG | 3.4 | - |
| ASN | 3.4 | -9.68 | LYS | 3.4 | - |
| GLN | 3.4 | -9.38 | ASP | 3.4 | - |
| MET | 3.4 | -1.48 | GLU | 3.4 | - |
| CYS | 3.4 | -1.24 | PRO | - | - |

b

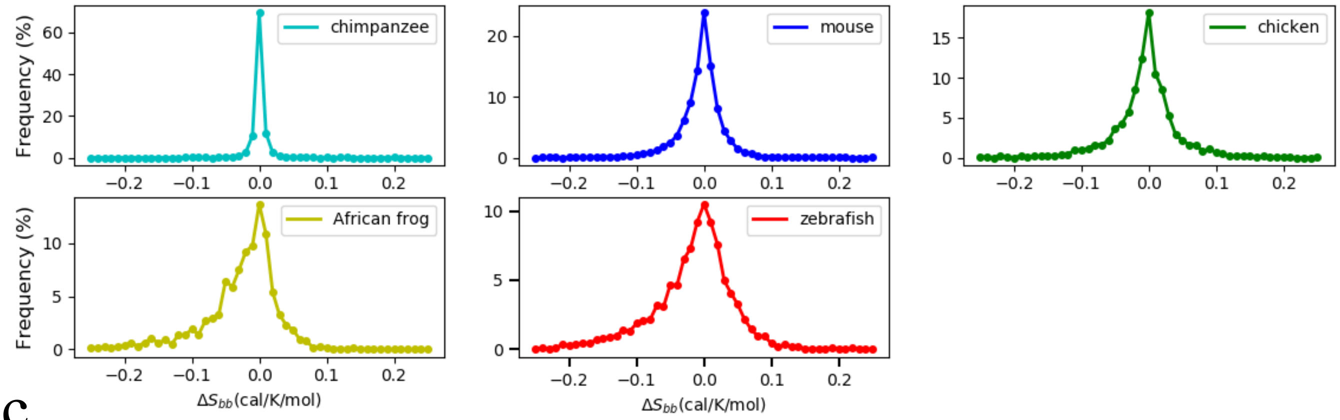

c

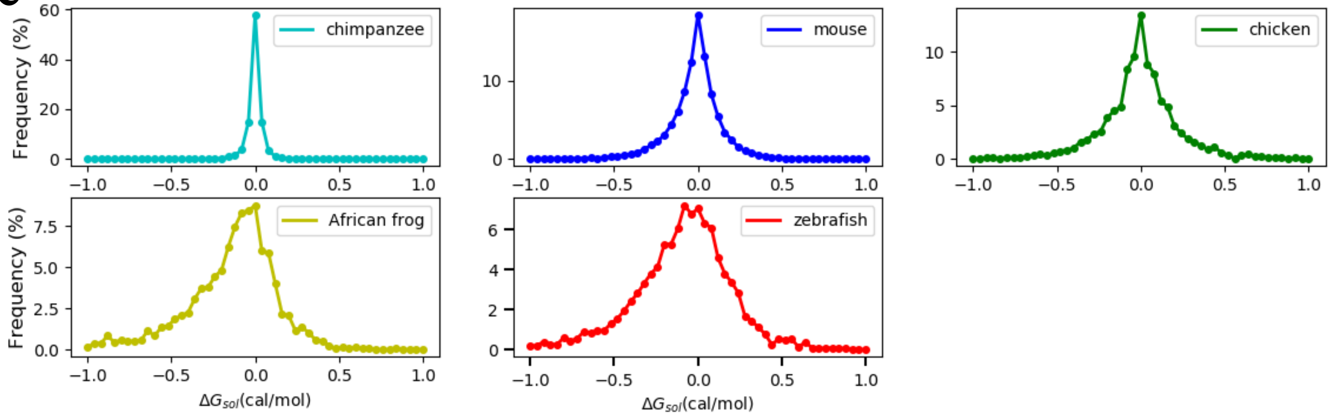

Table S4: A list of human proteins and the PDB identification codes used to build their initial structures for simulation in the structure-based energetics analysis.

| Protein Name | Templates |
|---|---|
| Acetylcholinesterase | 1f8u A |
| Actin-histidine N-methyltransferase | 6ox0 A |
| Actin, cytoplasmic 1 | 6anu F |
| Actin, cytoplasmic 2 | 5jlh A |
| Adenosine deaminase | 3iar A |
| Adenylate kinase 2, mitochondrial | 2c9y A |
| ADP-ribose glycohydrolase ARH3 | 6d36 A |
| ADP-ribosylation factor-like protein 2-binding protein | 3doe B |
| ADP-ribosylation factor-like protein 6 | 2h57 A |
| Anoctamin-10 | 5oc9 A |
| Aromatase | 3eqm A |
| Aspartoacylase | 2o4h A |
| ATP-dependent RNA helicase SUPV3L1, mitochondrial | 3rc3 A |
| Autophagy-related protein 101 | 4wzg A |
| Barrier-to-autointegration factor | 1ci4 A |
| Beta-2-microglobulin | 1a1m B |
| Betaine--homocysteine S-methyltransferase 1 | 1lt7 A |
| Bifunctional arginine demethylase and lysyl-hydroxylase JMJD6 | 6gdy A |
| BRISC and BRCA1-A complex member 2 | 6h3c C |
| BRO1 domain-containing protein BROX | 3uly A |
| C5a anaphylatoxin chemotactic receptor 1 | 5o9h A |
| Carbonyl reductase family member 4 | 4cql B |
| Carboxypeptidase O | 5mrv A |
| CCR4-NOT transcription complex subunit 7 | 4gmj B |
| CCR4-NOT transcription complex subunit 9 | 4cru B |
| Ceramide-1-phosphate transfer protein | 4k80 A |
| Chitinase domain-containing protein 1 | 3bxw A |
| Citrate synthase, mitochondrial | 5uzq A |
| Cleavage and polyadenylation specificity factor subunit 5 | 3mdg A |
| COP9 signalosome complex subunit 3 | 4d10 C |
| Cyclin-dependent kinase-like 1 | 4agu A |
| Cysteine dioxygenase type 1 | 2ic1 A |
| Cytochrome c oxidase subunit 1 | 5z62 A |
| Cytochrome c oxidase subunit 2 | 5z62 B |
| Cytochrome c oxidase subunit 3 | 5z62 C |
| Cytosolic phospholipase A2 | 1cjy A |
| D-aminoacyl-tRNA deacylase 1 | 2okv A |
| Dihydropyrimidinase-related protein 3 | 4bkn A |
| Diphosphomevalonate decarboxylase | 3d4j A |
| DNA polymerase beta | 1bpx A |
| DNA-directed RNA polymerase II subunit RPB7 | 2c35 B |
| Dynactin subunit 6 | 3tv0 A |
| Dynein assembly factor with WDR repeat domains 1 | 5nnz A |
| EKC/KEOPS complex subunit TPRKB | 3enp A |
| Endoribonuclease LACTB2 | 4ad9 A |

| | |
|---|---|
| Enhancer of rudimentary homolog | 1w9g A |
| Enolase-phosphatase E1 | 1yns A |
| Enoyl-[acyl-carrier-protein] reductase, mitochondrial | 2vcy A |
| Enoyl-CoA hydratase domain-containing protein 3, mitochondrial | 2vx2 A |
| Eukaryotic translation initiation factor 2 subunit 3 | 6k71 P |
| Exosome complex component MTR3 | 2nn6 F |
| Flap endonuclease 1 | 1ul1 X |
| Fructose-bisphosphate aldolase B | 1qo5 A |
| Fumarate hydratase, mitochondrial | 5d6b A |
| G1/S-specific cyclin-D1 | 2w96 A |
| Galectin-related protein | 2jj6 A |
| Glutathione S-transferase LANCL1 | 3e6u A |
| Glycerol-3-phosphate dehydrogenase 1-like protein | 2pla A |
| Glycine amidinotransferase, mitochondrial | 1jdw A |
| Glycolipid transfer protein | 1swx A |
| Glypican-1 | 4ywt A |
| GTPase NRas | 3con A |
| Guanidinoacetate N-methyltransferase | 3orh A |
| Guanine nucleotide-binding protein G(I)/G(S)/G(T) subunit beta-1 | 4pnk B |
| Guanine nucleotide-binding protein subunit beta-2-like 1 | 4aow A |
| Guanosine-3',5'-bis(diphosphate) 3'-pyrophosphohydrolase MESH1 | 3nr1 A |
| Haloacid dehalogenase-like hydrolase domain-containing protein 2 | 3hlt A |
| Haloacid dehalogenase-like hydrolase domain-containing protein 3 | 3k1z A |
| Hemoglobin subunit alpha | 1a00 A |
| Histamine N-methyltransferase | 1jqe A |
| Histone deacetylase 3 | 4a69 A |
| Histone deacetylase 8 | 1t64 A |
| Histone H3.3 | 3av2 A |
| Histone-binding protein RBBP4 | 3gfc A |
| Histone-binding protein RBBP7 | 3cfs B |
| Hypoxia-inducible factor 1-alpha inhibitor | 1h2k A |
| Inositol-tetrakisphosphate 1-kinase | 2qb5 A |
| L-lactate dehydrogenase A chain | 1i10 A |
| Lanosterol 14-alpha demethylase | 3jus A |
| LYR motif-containing protein 4 | 5wgb B |
| Malignant T-cell-amplified sequence 1 | 5ons A |
| Methionine adenosyltransferase 2 subunit beta | 2ydy A |
| Methionine aminopeptidase 1 | 4iu6 A |
| Methylthioribose-1-phosphate isomerase | 4ldq A |
| Mitogen-activated protein kinase 12 | 1cm8 A |
| Mitotic spindle assembly checkpoint protein MAD2B | 3abd A |
| mRNA export factor | 3mmy A |
| Mth938 domain-containing protein | 2ab1 A |
| Myoglobin | 3rgk A |
| Myotrophin | 3aaa C |
| Myotubularin-related protein 2 | 1lw3 A |
| N-acetylneuraminate lyase | 6arh A |
| N-alpha-acetyltransferase 50 | 2ob0 A |
| N-terminal Xaa-Pro-Lys N-methyltransferase 1 | 5cvd A |

| | |
|---|---|
| NAD-dependent protein deacetylase sirtuin-2 | 1j8f A |
| NAD-dependent protein deacylase sirtuin-5, mitochondrial | 2b4y A |
| NADH-ubiquinone oxidoreductase chain 1 | 5xtc s |
| NADH-ubiquinone oxidoreductase chain 2 | 5xtc i |
| NEDD8-activating enzyme E1 regulatory subunit | 1tt5 A |
| NEDD8-conjugating enzyme UBE2F | 3fn1 B |
| Neutral ceramidase | 4wgk A |
| Nicotinate phosphoribosyltransferase | 4yub A |
| Nuclear cap-binding protein subunit 2 | 1h2t Z |
| O-phosphoseryl-tRNA(Sec) selenium transferase | 4zdl A |
| Obg-like ATPase 1 | 2ohf A |
| Osteoclast-stimulating factor 1 | 3ehq A |
| Pachytene checkpoint protein 2 homolog | 5vqa A |
| Peroxisomal 2,4-dienoyl-CoA reductase | 4fc6 A |
| Phospholysine phosphohistidine inorganic pyrophosphate phosphatase | 2x4d A |
| Pre-mRNA-splicing factor 38A | 5o9z I |
| Probable tRNA N6-adenosine threonylcarbamoyltransferase | 6gwj K |
| Proliferating cell nuclear antigen | 1axc A |
| Protein argonaute-3 | 5vm9 A |
| Protein cereblon | 6bn7 B |
| Protein mago nashi homolog | 2hyi A |
| Protein MEMO1 | 3bcz A |
| Protein N-lysine methyltransferase METTL21A | 4lec A |
| Protein phosphatase 1 regulatory subunit 7 | 6hkw A |
| Protein rogdi homolog | 5xqi A |
| Protein-L-isoaspartate(D-aspartate) O-methyltransferase | 1i1n A |
| Protein-tyrosine sulfotransferase 1 | 5wri A |
| Protein/nucleic acid deglycase DJ-1 | 1j42 A |
| Proto-oncogene tyrosine-protein kinase Src | 2h8h A |
| Putative deoxyribonuclease TATDN1 | 2xio A |
| Putative nucleotidyltransferase MAB21L1 | 5eom A |
| Queuine tRNA-ribosyltransferase catalytic subunit 1 | 6h42 A |
| Ran guanine nucleotide release factor | 5yfg A |
| Ras-related protein Rab-4A | 2bmd A |
| Ras-related protein Rab-4B | 2o52 A |
| Ras-related protein Rap-1b | 4dha A |
| Regucalcin | 4gnb A |
| Ribonucleoside-diphosphate reductase large subunit | 3hnc A |
| Ribonucleoside-diphosphate reductase subunit M2 | 2uw2 A |
| RNA polymerase II subunit A C-terminal domain phosphatase SSU72 | 3o2s B |
| RNA-binding protein PNO1 | 6g18 x |
| S-methyl-5'-thioadenosine phosphorylase | 1cb0 A |
| Selenide, water dikinase 1 | 3fd5 A |
| SH3 domain-binding glutamic acid-rich-like protein 2 | 2ct6 A |
| Sigma non-opioid intracellular receptor 1 | 5hk1 A |
| Small nuclear ribonucleoprotein E | 4f7u E |
| Sortilin | 3f6k A |

| | |
|---|---|
| SOSS complex subunit C | 4owt C |
| Superoxide dismutase [Cu-Zn] | 1azv A |
| TBC1 domain family member 7 | 3qwl A |
| Thioredoxin domain-containing protein 17 | 1wou A |
| Transcription elongation factor SPT4 | 3h7h A |
| Transmembrane protein 14C | 2los A |
| Ubiquitin carboxyl-terminal hydrolase 46 | 5cvm A |
| Ubiquitin-conjugating enzyme E2 S | 5l9t T |
| Ubiquitin-conjugating enzyme E2 T | 4ccg A |
| Ubiquitin-conjugating enzyme E2 variant 2 | 1j74 A |
| Ubiquitin-fold modifier-conjugating enzyme 1 | 2z6o A |
| Ubiquitin-like protein 5 | 4pyu A |
| UMP-CMP kinase | 1tev A |
| Uroporphyrinogen decarboxylase | 1jph A |
| Vacuolar protein sorting-associated protein 26A | 2fau A |
| Vacuolar protein sorting-associated protein 29 | 1w24 A |
| WD repeat domain phosphoinositide-interacting protein 3 | 6iyy A |
| WD repeat-containing protein 48 | 5cvo B |
| WD repeat-containing protein 61 | 3ow8 A |
| WD repeat-containing protein 92 | 3i2n A |
| 3-hydroxyanthranilate 3,4-dioxygenase | 5tk5 A |
| 39S ribosomal protein L41, mitochondrial | 3j7y 9 |
| 4-hydroxy-2-oxoglutarate aldolase, mitochondrial | 3s5n A |
| 4-hydroxyphenylpyruvate dioxygenase | 5ec3 A |
| 40S ribosomal protein S15a | 5a2q W |
| 40S ribosomal protein S3a | 5a2q B |
| 60S ribosome subunit biogenesis protein NIP7 homolog | 1sqw A |

Figure S4: Cold adaptation of L-Lactate dehydrogenase A chain (LDHA) of Antarctic fishes in extreme cold. (a) Molecular simulations of LDHA proteins for three difference organisms at three different temperatures. The table lists values for each of the two independent simulations. Each simulation is 500 ns in length. RMSD values for each simulation are listed. (b) RMSF for LDHA of different species at temperatures 37 ℃, 17 ℃ and 0 ℃ - left to right column.

a

| Simulation System | T=37 ℃ RMSD (Å) | T=17 ℃ RMSD (Å) | T=0 ℃ RMSD (Å) |
|---|---|---|---|
| Mackerel icefish | 3.51±0.30<br>2.16±0.24 | 3.66±0.27<br>2.23±0.18 | 2.10±0.31<br>3.71±0.54 |
| Common carp | 3.42±0.34<br>3.06±0.50 | 2.10±0.24<br>2.76±0.27 | 2.40±0.22<br>2.34±0.49 |
| Human | 2.59±0.45<br>2.73±0.23 | 1.79±0.20<br>1.70±0.14 | 1.88±0.23<br>1.85±0.31 |

b

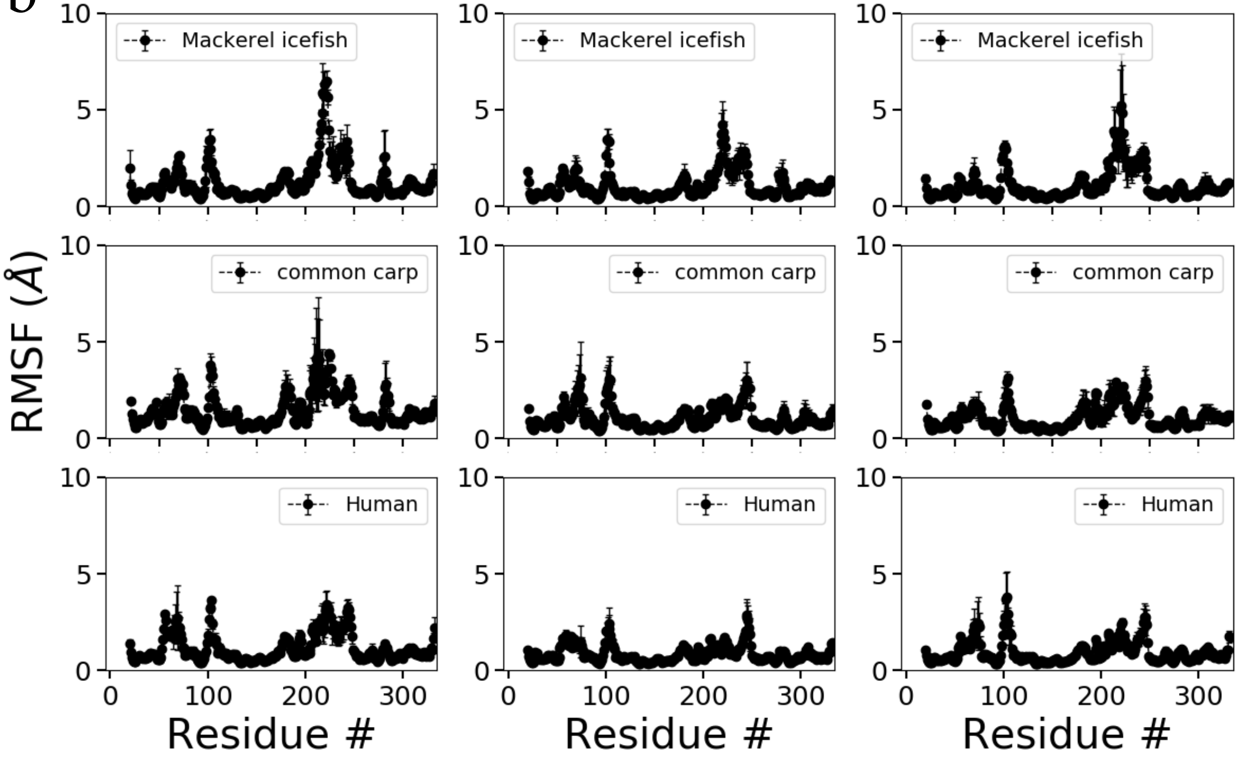

Figure S5: Cold adaptation of L-Lactate dehydrogenase A chain (LDHA) of Antarctic fishes in extreme cold. (a) Clustering for LDHA conformations in the simulation of the protein from three different organisms at three different temperatures (37 ℃, 17 ℃, 0 ℃). (b) Sequence alignment between Mackerel icefish, Common carp and Human.

a

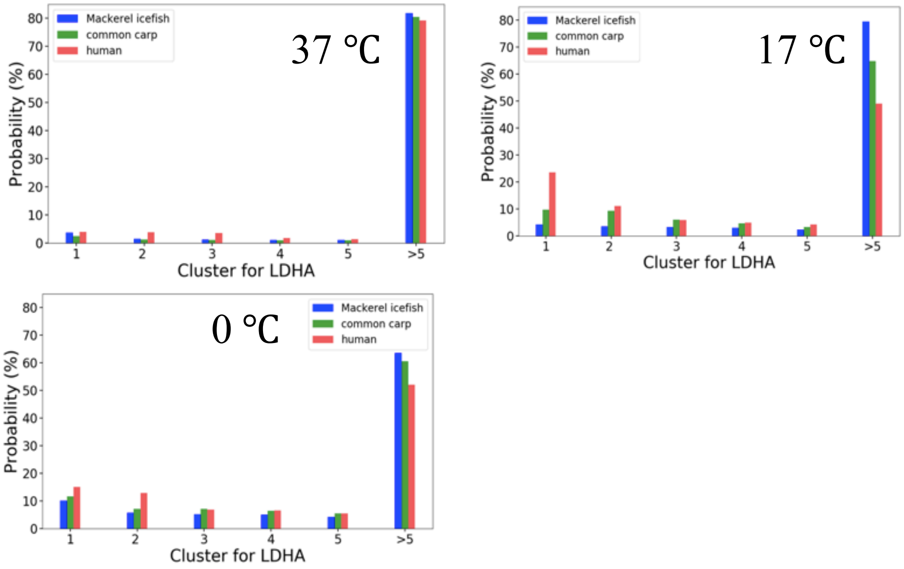

b

```
M-STKEKLIS HVMKEEPVGS RSKVTVVGVG MVGMASAISI LLKDLCDELA MVDVMEDKLK     59
MASTKEKLIT HVSKEEPAGP TNKVTVVGVG MVGMAAAISI LLKDLTDELA LVDVMEDKLK     60
MATLKDQLIY NLLKEE-QTP QNKITVVGVG AVGMACAISI LMKDLADELA LVDVIEDKLK     59
* : *::**  :: ***      .*:******  ****.**** *:*** **** :***:*****

GEVMDLQHGS LFLKT-KIVG DKDYSVTANS KVVVVTAGAR QQEGESRLNL VQRNVNIFKF     118
GEAMDLQHGS LFLKTHKIVA DKDYSVTANS KVVVVTAGAR QQEGESRLNL VQRNVNIFKF     120
GEMMDLQHGS LFLRTPKIVS GKDYNVTANS KLVIITAGAR QQEGESRLNL VQRNVNIFKF     119
** ******* ***:* ***. .***.***** *:*::***** ********** **********

IIPNIVKYSP NCILMVVSNP VDILTYVAWK LSGFPRHRVI GSGTNLDSAR FRHLIGEKLH     178
IIPNIIKYSP NCILLVVSNP VDILTYVAWK LSGLPRNRVI GSGTNLDSAR FRHLMGEKLG     180
IIPNVVKYSP NCKLLIVSNP VDILTYVAWK ISGFPKNRVI GSGCNLDSAR FRYLMGERLG     179
****::**** ** *::**** ********** :**:*::*** *** ****** **:*:**:*

LHPSSCHAWI VGEHGDSSVP VWSGVNVAGV SLQGLNPQMG TEGDGENWKA IHKEVVDGAY     238
IHPSNCHGWV IGEHGDSSVP VWSGVNVAGV FLQGLNPDMG TDKDKEDWKS VHKMVVDSAY     240
VHPLSCHGWV LGEHGDSSVP VWSGMNVAGV SLKTLHPDLG TDKDKEQWKE VHKQVVESAY     239
:** .**.*: :********* ****:*****  *: *:*::* *: * *:**  :** **:.**

EVIKLKGYTS WAIGMSVADL VESIIKNMHK VHPVSTLVQG MHGVKDEVFL SVPCVLGNSG     298
EVIKLKGYTS WAIGMSAADL CQSILKNLRK CHPVSTLVKG MHGVNEEVFL SVPCILGNSG     300
EVIKLKGYTS WAIGLSVADL AESIMKNLRR VHPVSTMIKG LYGIKDDVFL SVPCILGQNG     299
********** ****:*.***  :**:**:::  *****:::* ::*::::*** ****:**:.*

LTDVIHMTLK AEEEKQLQKS AETLWGVQKE LTL      331
LTDVVHMTLK SDEEKQLVKS AETLWGVQKD LTL      333
ISDLVKVTLT SEEEARLKKS ADTLWGIQKE LQF      332
::*::::**. ::** :* ** *:****:**: * :
```
